# Metric Ion Classification (MIC): A deep learning tool for assigning ions and waters in cryo-EM and x-ray crystallography structures

**DOI:** 10.1101/2024.03.18.585639

**Authors:** Laura Shub, Wenjin Liu, Georgios Skiniotis, Michael J. Keiser, Michael J. Robertson

## Abstract

At sufficiently high resolution, x-ray crystallography and cryogenic electron microscopy are capable of resolving small spherical map features corresponding to either water or ions. Correct classification of these sites provides crucial insight for understanding structure and function as well as guiding downstream design tasks, including structure-based drug discovery and de novo biomolecule design. However, direct identification of these sites from experimental data can prove extremely challenging, and existing empirical approaches leveraging the local environment can only characterize limited ion types. We present a novel representation of chemical environments using interaction fingerprints and develop a machine-learning model to predict the identity of input water and ion sites. We validate the method, named Metric Ion Classification (MIC), on a wide variety of biomolecular examples to demonstrate its utility, identifying many probable mismodeled ions deposited in the PDB. Finally, we collect all steps of this approach into an easy-to-use open-source package that can integrate with existing structure determination pipelines.

## Introduction

Hydration and ion binding are vital for biomolecular function, contributing to structure,^1,2^ ligand binding,^3–5^ enzymatic catalysis,^6^ and dynamics.^1,7^ Proper rationalization of their effects in structures requires accurate identification of these sites as either water or a specific ion bound. However, pinpointing the identity of spherical features in experimental maps can be challenging, as the experimental data may be of insufficient quality or interpretability to definitively classify the scatterer. In structures derived from x-ray crystallography, individual ions and water can often be distinguished by examining the Fo-Fc difference map or OMIT map.^8^ However, this depends on data quality and can be difficult if the scattering is similar, for example when differentiating between water, sodium, and magnesium. Cryogenic electron microscopy data is even more problematic due to intrinsic challenges in generating meaningful difference maps.^9^ While scattering differences between atoms of different charges in certain resolution ranges can be used to discriminate atomic charges from cryo-EM data in theory,^10^ this often proves difficult in practice.

Rather than directly determining the identity of the spherical feature from the experimental data, one can also consider the environment around the feature that is responsible for its coordination. Cations generally have a coordination shell of several partial or formal negatively charged atoms with well defined geometry and coordination distances.^11^ Water has an ideal tetrahedral coordination with two hydrogen bond donors and two hydrogen bond acceptors. Anions tend to have a less well-defined coordination shell than cations but with positive interaction partners, specifically the guanidinium of arginine.^12^ While computational tools exist for classification based on the local environment, these methods tend to be focused on more specific subsets, for example comparing different metal ions^13–16^ or validating waters.^17^

In recent years, machine learning (ML) has been successfully applied to various biomolecular modeling tasks including structure prediction,^18,19^ protein design,^20^ molecular docking,^21,22^ and molecular dynamics simulations.^23^ The field of cryo-EM specifically has seen increased interest in ML for modeling proteins^24–27^ and nucleic acids^28^, as well as improving and evaluating map and model quality.^29–31^ ML methods have been explored for identifying potential binding sites for specific ions or ion subclasses,^32–34^ but assigning the identity of experimentally determined sites remains underexplored. One complicating factor is the relative scarcity of high-quality experimental structures with the full set of possible ions bound compared to what is needed for the 3-dimensional convolutional architectures typically used in these applications, highlighting the need to consider both alternate site representations and model architectures for this task.

In cheminformatics, molecular fingerprints are vector representations that encode chemical structure.^35,36^ They have been used in mapping chemical space,^37^ virtual drug screening,^38^ and as input to quantitative structure-activity relationship models.^39^ In recent years, this concept has been expanded to interface fingerprints that capture the geometry of ligand-receptor complexes.^40–43^ These representations have been used to filter virtual screening results by binding mode and as inputs to ML models for binding affinity prediction.^44,45^ One such example is the extended interaction fingerprint presented as part of the LUNA Python toolkit,^46^ designed for calculating and encoding protein-ligand interactions. In this work, we implement an extension to LUNA to generate identity-blinded geometric fingerprints that capture the chemical microenvironment of ion and water coordination sites.^47^

Deep metric learning is an ML framework employed for facial recognition, anomaly detection, and signature verification applications.^48–51^ In contrast to classic ML approaches trained to predict a 1:1 label for each example, deep metric learning models learn an efficient virtual landscape that maximizes the distance between objects of different classes. It has been ues in cheminformatics to learn molecular similarity and improve molecular property prediction.^52,53^ Metric models are often trained on triplets of examples sampled from each batch, consisting of an anchor, a positive example from the same class as the anchor, and a negative example of a different class.^54–56^ While this near-cubic increase in potential training examples can introduce implementation challenges, it is a useful property when operating in a data-sparse regime. We exploit this property by training a metric learning model on a relatively small dataset of ions in structures from the Protein Data Bank (PDB) to learn a low-dimensional embedding (landscape) conditioned on deposited ion identity, and use these embeddings for downstream identity classification.

Here, we present Metric Ion Classifier (MIC), an open-source tool for assigning identities to sites in PDB structures. MIC utilizes a novel ion-fingerprint representation and deep metric learning approach to predict the class of placed ions and waters in input structures. We demonstrate MIC’s accuracy on a test set of structures from the PDB and use explainable AI feature attribution techniques to understand the biophysical rationale behind these predictions. Finally, we evaluate the performance on diverse x-ray crystallography and cryo-EM structures of both proteins and RNA, demonstrating the widespread utility of this approach. We hope this will prove a vital verification tool in structural biology workflows and represent an important step towards interpretable machine learning in the field.

## Results

### Architecture and Performance of MIC

The MIC tool assigns identities to waters and ions modeled in PDB structures. The overall workflow consists of three steps: 1) generating the fingerprint representation for the chemical environment of the density, 2) condensing this representation into a lower-dimensional embedding using a trained deep metric model, and 3) passing this embedding through a support vector classifier (SVC) to obtain final probabilities for all classes as well as prediction confidence **(Fig. 1)**. The class with the highest probability is then taken as the final prediction.

**Fig. 1:**
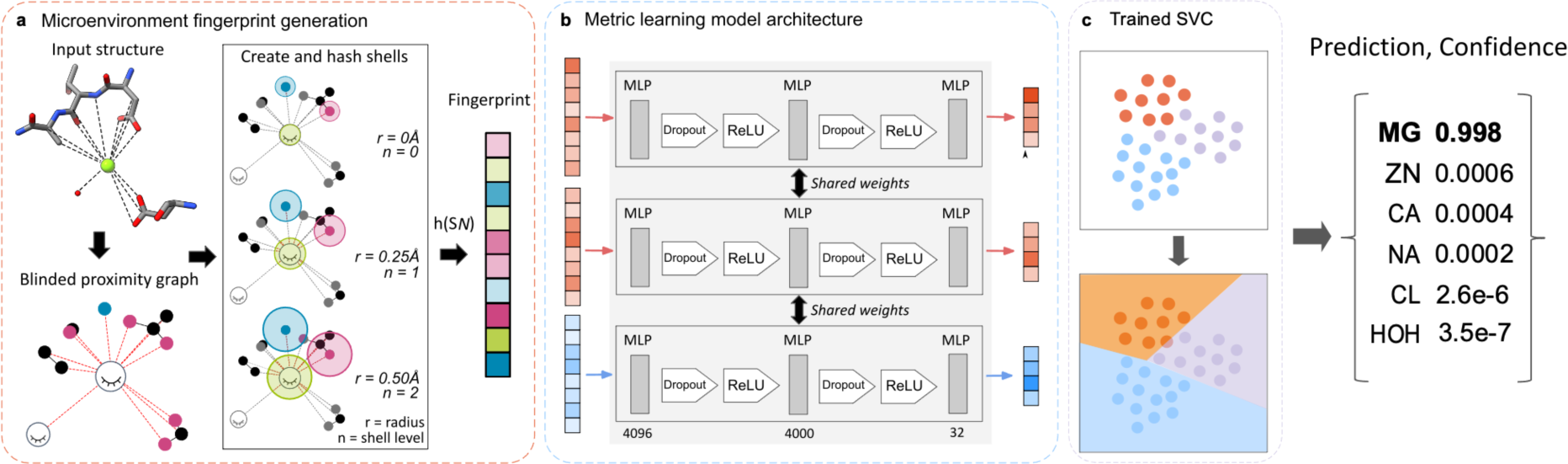
Overview of MIC workflow. MIC is a multi-step ML workflow for classifying experimental water and ion sites. **a**, Ion fingerprints are generated by first constructing a proximal interaction graph containing all atoms within 6A for the density of interest. The fingerprint generation protocol iteratively captures local chemical information by hashing the atomic invariants and interactions within consecutive shells originating from each atom. The example structure shown here is 4KU4:A:Mg:302. **b**, The fingerprints are embedded into a lower dimensional embedding space by a metric learning model consisting of a 4096-dimensional input layer, a single hidden layer with 4000 neurons, and an output layer of 32. ReLU is an activation function *x* = *max*(*x*, 0). **c**, The final step of MIC is using an SVC on the generated fingerprints to output probabilities for each class. The class with the highest probability is taken as the predicted label.

Prior approaches to conceptually similar tasks have used voxel representations as input to neural networks, necessitating large 3D-convolutional architectures that are both orientation-dependent and rely on abundant training data to tune properly.^33,34^ We overcome these limitations in two ways. To represent each density, we use a modified version of the LUNA toolkit developed to calculate intermolecular interactions at protein-ligand interfaces and encode them into a fixed-length vector representation known as a “fingerprint”. This greatly reduces the size of our models while meaningfully capturing information required for downstream classification. The generation process for these ion fingerprints is closely related to interface fingerprints with a few key exceptions **(Fig. 1a)**. First, a proximity graph is constructed comprising all atoms < 6Å from the center of mass of the ion or water of interest. Each atom in this graph is assigned a set of atomic identifiers depending on its chemical identity and user-selected fingerprint type **(Extended Data Table 1).** Crucially, we remove the initial features of both the density itself and any additional waters or ions in the graph, effectively blinding the representation to any existing label to protect against data leakage during downstream prediction. The final list of modified atomic identifiers and all interactions are passed through a distance-dependent hash function that converts these input features into numeric values, which are folded down to 4,096 dimensions following standard molecular fingerprinting procedures.^35,47^

These fingerprints are further condensed using a deep metric model. This model, constructed as a small feed-forward network, is trained to learn low-dimensional embeddings that maximize the distance between members of different classes (**Fig. 1b**). This step establishes the discriminative capabilities of the model, enabling accurate differentiation between closely related density types. The final predictions are generated by an SVC that uses these learned latent embeddings to calculate a probability for each class, the maximum of which is taken as the label (**Fig. 1c**). Full details for fingerprint generation and model training are provided in Methods, and the full list of sites used for training and testing are provided in **Supplementary Table 1** and **Extended Data Fig. 1a**. The model was trained for 1000 epochs (**Extended Data Fig. 2a,b**).

In **Figure 2**, we present the performance of the MIC protocol trained on the six most prevalent classes from our curation of the Protein Data Bank: water, magnesium, sodium, zinc, calcium, and chloride. The model achieves an initial accuracy of 76.7% on a held-out test set and displays notable trends in performance by class, specifically showing high accuracy for zinc, magnesium, calcium, and water recovery. (**Fig. 2c, Supplementary Table 2**). A particularly interesting property of these learned embeddings is the organization by charge, visualized here with UMAP and confirmed by both PCA and the learned latent embeddings (**Fig 2a,b, Extended Data Fig. 2c-f**). This constraint was not explicitly included in the representation or loss function during training, and the model was provided with no information about class relationships. Rather, this is an emergent learned quality of the transition of the chemical microenvironment of the sites themselves. The model’s ability to learn the underlying structure inherent to the dataset supports the utility of our representation in capturing relevant information. Additionally, this reasonably organized continuous landscape also allows for confidence estimation through proximity to the classifier decision boundary, discussed below.

**Fig. 2:**
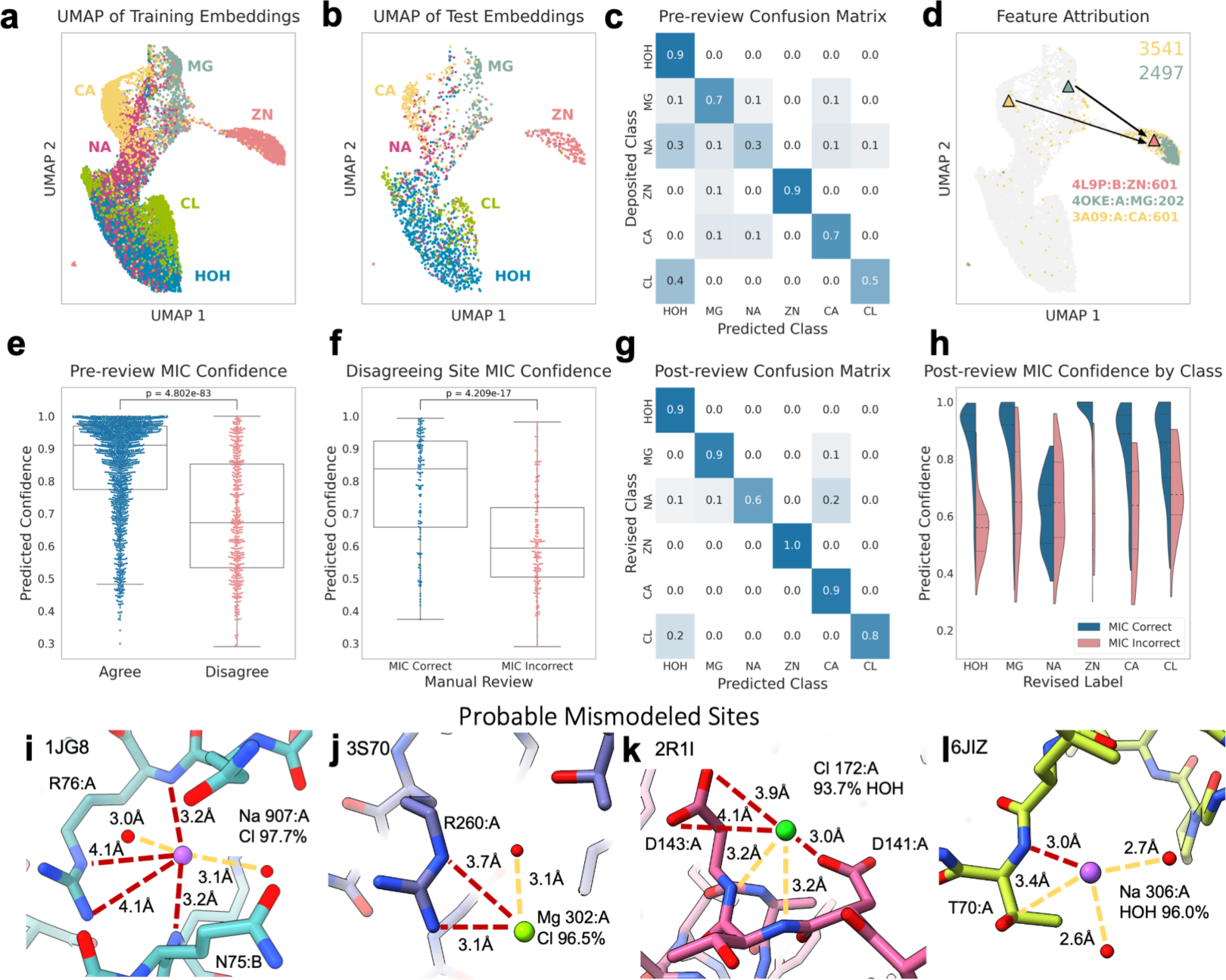
MIC learned embeddings, performance, and validation. **a-b**, UMAP visualization of training and test set embeddings from the MIC prevalent-set model, colored by deposited class. **c**, Confusion matrix of the deposited labels and MIC predicted labels for the test set. **d,** UMAP visualization of training set embeddings, colored by the value of the bits 2497 (green) and 3541 (yellow), corresponding to the presence of a cysteine sulfur and imidazole nitrogen, respectively. The triangles indicate the position of specific examples used to perform feature attribution:4OKE:A:Mg:202 (green), 3A09:A:Ca:601 (yellow), and 4L9P:B:Zn:601 (pink). **e,** Comparison of the confidence values for MIC predictions that agree vs disagree with the deposited label. **f,** Comparison of confidence values for manually inspected disagreeing examples with accurate vs inaccurate MIC-predicted labels. **g,** Confusion matrix of revised labels and MIC predictions following manual review of disagreeing test examples. **h**, Violin plots of the confidence of correct vs incorrect MIC test set predictions, split by class. **i-l,** Examples of disagreeing annotations with probable mismodeling. **i**, Sodium in 1J86 corrected to a chloride, 97.7% confidence. **j**, Magnesium in 3S70 corrected to chloride, 96.5% confidence. **k**, Chloride in 2RL1 corrected to water, 93.7% confidence. **l**, Sodium in 6JIZ corrected to water, 96.0% confidence. Red dashed lines depict unfavorable interactions in the originally modeled structure.

One potential drawback to MIC, and in fact most machine learning-based approaches, is a lack of interpretability of the resulting models, also known as the “black box” problem. We aimed to address this and provide further validation of the model through pairwise feature attribution with integrated gradients, a technique used to quantify the importance of input features to the model’s output.^57,58^ By calculating the attribution of fingerprints near the centroid of an ion cluster in embedding space, we can form hypotheses about which bits in the input fingerprint are most salient for a given class. Furthermore, we can use LUNA to trace back these features to their origin in the input structure’s atoms or interactions, allowing us to support the predictions with a biophysical rationale (The full details of feature attribution protocol as implemented by L. Ponzoni, PhD are provided in the Methods section).

To investigate the model’s rationale behind the emergent organization by chemical microenvironment in the embedding space, we used pairwise attribution to probe the features most useful to the model for differentiating between closely related classes. Comparing two representative zinc and magnesium fingerprints (4L9P:B:ZN:601 and 4OKE:A:MG:202, **Extended Data Fig. 2j,k**) provides insight into how the model separates these embeddings despite similar charges. The nearby Cys367 sulfur appears in the top features by importance for zinc when compared against magnesium along with the short distance to the Asp365 sidechain carboxylate group (**Supplementary Table 3**). Visualizing the embedding space by the value of the corresponding fingerprint bit (2497) shows strong localization in the zinc cluster, following known properties of zinc binding sites and likely contributing to the high confidence prediction for this example **(Fig. 2d**).^59^ Conversely, our analysis showed that salient features for magnesium similarly prioritized the slightly longer distances of nearby carboxylates (Asp6, Glu8) and the number of nearby waters, commonly observed features of magnesium sites.^60^ Comparing this same zinc against a calcium example (3BMV:CA:A:684, **Extended Data Fig. 2l**) also yields known important features. In addition to the Cys367 and Asp365 recovered against magnesium, the attribution value of bit 3541 corresponding to the nearby His433 imidazole nitrogen is higher, indicating additional importance of this feature for the model in distinguishing calcium and zinc examples (**Fig. 2d**). In comparing the calcium and magnesium fingerprints, both assign high attribution to oxygens in their top features, but calcium includes backbone carbonyl oxygens while magnesium again includes the Asp6 carboxyl and the AMP phosphate oxygen, agreeing with known properties of these sites.^60,61^ Interestingly, one feature that consistently returned high attribution was the null shell corresponding to the ion itself and any proximal waters and ions. The magnitude of the value at this index is consistently high and indicates the number of total sites encoded in the representation, a feature that is evidently useful to the model in structuring the learned latent space.

In addition to predicting the identity of a site, MIC provides a measure of confidence through the probability estimates output by the SVC. Because the latent representation transitions smoothly between chemically related classes, we can use the proximity to the decision boundary to measure confidence in a given prediction. Indeed, we found that our model was well-calibrated such that this simple metric showed statistically significant separation between test set predictions that agreed and disagreed with the deposited PBD label (P-value<<<1e^-10^, **Fig. 2e, Extended Data Table 3**). This property could assist the user in interpreting MIC results and encouraged us to further investigate these high-confidence disagreeing examples from the test set.

### Manual Inspection of Disagreeing Sites

Following model prediction, we manually reviewed 455 disagreeing test examples and considered what the correct label should be based upon several factors including favorable/unfavorable interactions, experimental map agreement (x-ray structures were re-refined with the alternative density and Fo-Fc maps were inspected in both cases), and coordination geometric features (**Extended Data Fig. 3a, Supplementary Table 2**). We assigned each structure a score between -3 and 3, with increasingly positive scores denoting more support for the MIC prediction and increasingly negative scores support for the original label. We identified 135 sites where we believe the provided label in the PDB to be incorrect and MIC accurate in its assignment and 176 sites where MIC is likely incorrect and the deposited label is correct. A further 80 sites were scored as 0, reflecting that even after manual inspection and re-refinement it was unclear which of the two labels were correct. 64 sites were also labeled as having unusual issues that would prevent proper prediction, including extended densities indicating the site represents a larger chemical entity than an ion or water, extensive heterogeneity and/or partial occupancy, the presence of an unusual multi-ion cluster, or that the likely correct identity of the ion did not fall within the set of predicted ions; indicated by a manual label of 30 (**Extended Data Fig. 3d,e**). In the manually annotated cases where the MIC assignment was correct over the deposited label, the average confidence was 78.0±17.2%, while the confirmed incorrect MIC predictions had an average confidence of 61.5±15.4% (**Fig. 2f**). The revised overall test set accuracy following manual annotation is 83.3% with an average confidence of 84.8±15.3% for correct predictions (**Fig. 2g**). The most common corrections made by MIC were reassigning spurious sodium and chloride ions to water (53 and 18 examples, respectively), followed by reassigning sodium to chloride and calcium to magnesium (11 examples) (**Extended Data Fig. 2b**). Given that 52 of the 258 total sodium sites in the test set were changed upon manual review, up to 20% of the sodium in the PDB may instead be water and up to 25% of all sodium in the PDB could be misannotated. Manually reviewing these examples additionally allowed us to provide an estimated accuracy cutoff by confidence (**Fig. 2h**). Except for sodium, the confidence of correctly predicted examples was significantly higher (P-value < 1e-5, **Extended Data Table 3**) than mispredicted examples. Overall, we found that a confidence of 70% was a useful cutoff in practice for most classes, and predictions below this cutoff typically require further review. Sodium is more challenging to predict confidently, likely due to the modest quality of annotated sodium ions in the dataset, and these predictions often require additional inspection.

Four diverse examples of high-confidence probable mismodeling captured by MIC are presented in **Fig. 2i-l**, showing a sodium to chloride (PDB:1JG8), magnesium to chloride (PDB:3S70), chloride to water (PDB:2RL1), and sodium to water (PDB:6JIZ) substitution. In each case, there is at least one short-range (3.0-3.2 Å) unfavorable interaction, and often several modest range (3.5-4.0 Å) unfavorable charge interactions, while lacking any opposite charge/partial charge interactions that would support the original assignment (for example, carbonyl interactions with a cation). None of the three deposited cations has the extended coordination shell or short coordination distances one would expect of a cation. Further, typically the experimental difference maps were improved upon re-refinement with the MIC ion (**Extended Data Fig. 3f-g)**, providing additional support for the corrected label.

### Validation of MIC on Structures Derived from Cryo-EM Maps

As all but 9 structures in the training set derive from x-ray crystallography, we wanted to examine how well MIC would work on cryo-EM structures. For this purpose we examined two disparate cases, representing the lower bound of resolution where an ion can still be resolved in a cryo-EM map (structures of melanocortin receptor 4 (MC4R) with bound calcium, nominal reported resolutions ranging from 2.6 Å to 3.1Å) and the upper bound of resolutions currently possible with cryo-EM (apoferritin, 1.15-1.27Å nominal resolution). In the first case, three different groups have determined several structures of MC4R bound to various ligands, resolving in each a spherical feature in the map thought by all three groups to be the calcium that has been biochemically demonstrated to be necessary for MC4R ligand binding.^62–65^ Further, some of the structures also resolve water molecules providing additional coordination for calcium ion binding. In the single structure from Israeli *et al.*^62^ of MC4R bound to setmelanotide (PDB:7AUE, **Fig. 3b**), MIC correctly identified calcium with 56.0% confidence, followed by sodium with 22.8% and magnesium with 13.8% confidence. In contrast, in the only other structure with an identical ligand, PDB:7PIU^63^ (**Fig. 3a**), the site was predicted to be either water (63.1%) or sodium (31.0%). This likely stems from the unexpectedly long carboxylate-calcium interaction distances modeled (**Fig. 3a**), which at 2.9-3.4Å are substantially longer than the ∼2.4Å average one would expect for a carboxylate-calcium interaction.^14^ These coordination distances are similar to those of the other structure from Heyder *et al.*, PDB:7PIV^63^ (**Fig. 3f**), which MIC predicted to be sodium (40.4%) or calcium (35.9%), with the improved classification likely due to the presence of an additional carbonyl interaction. All four structures from Zhang *et al.*^64^ (PDB:7F53, 7F54, 7F55, 7F58; **Fig. 3g,c-e**) are predicted to have a calcium ion at this site with high confidence (96.4%, 77.3%, 84.9%, 90.8%). Given the biochemical demonstration in Yu *et al.*^65^ that this is the site responsible for the calcium-dependence of ligand binding, all structures almost certainly did make the correct assignment as calcium, a result typically correctly predicted by MIC. In the case of 7PIV and particularly 7PIU, the discrepancy can be attributed to unusual coordination modeling, which is not unexpected in the ∼2.5-3.0Å nominal resolution range where ions can begin to be resolved but extremely precise placement of sidechain atoms remains challenging. Thus MIC in this resolution range also provides some level of audit on the overall modeling of the ion/water coordination site.

**Fig. 3:**
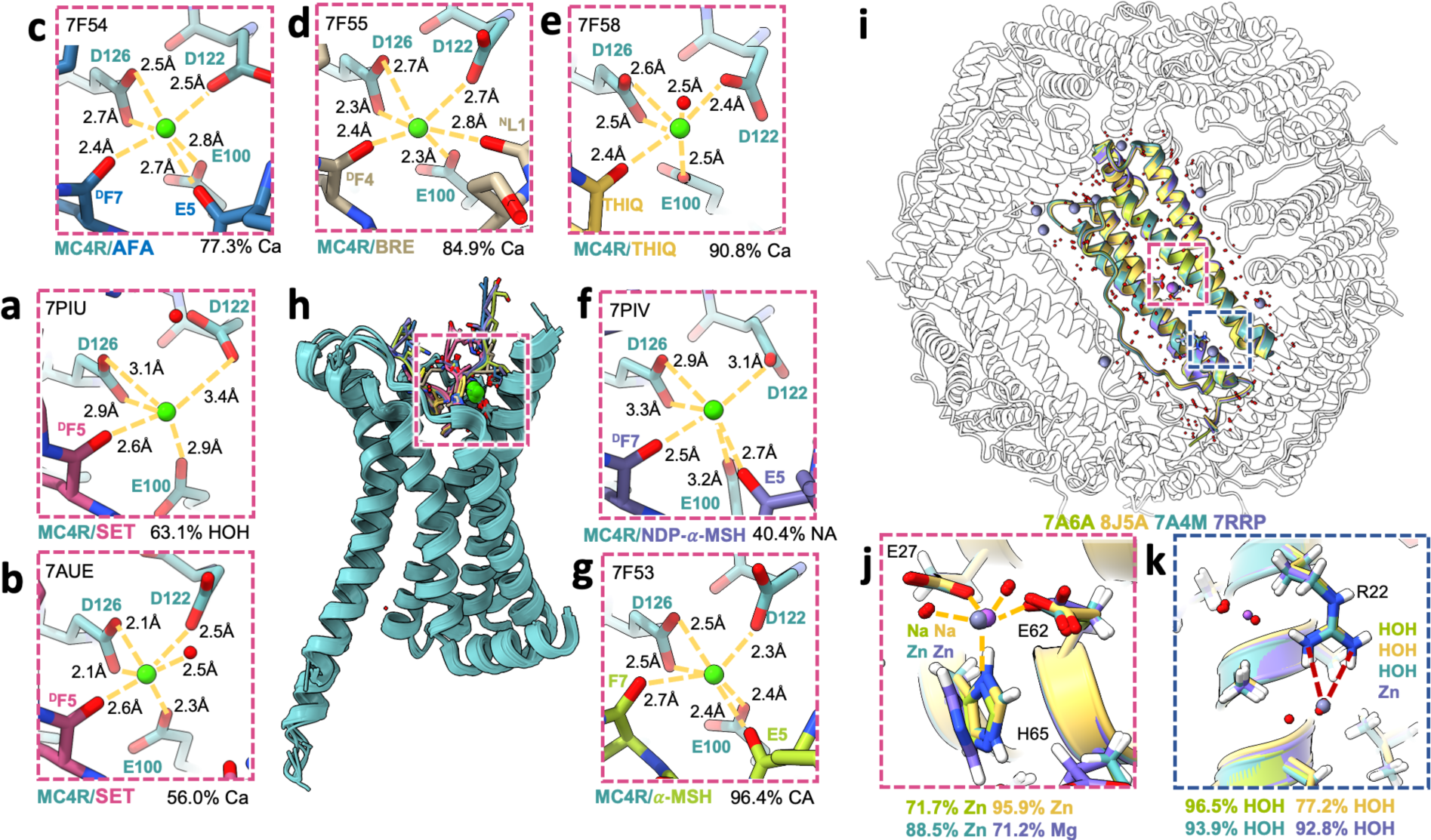
MIC predictions on Cryo-EM structures of MC4R and apoferritin. **a**-**h,** MC4R Ca^2+^-coordination site in complex with various ligands: setmelanotide (SET, **a,b**), afamelantodie (AFA, **c**), bremelanotide (BRE, **d),** THIQ (**e**), NDP-α-MSH (**f)**, and α-MSH (**g)**. **i-k,** Superimposed ion coordination sites in four apoferritin structures: 7A4M (green), 7RRP (purple), 7A6A (teal), 8J5A (yellow). **j,** For three structures, the ion is predicted to be zinc with confidence exceeding 70%. The 7RRP outwardly turned histidine imidazole shifts the prediction from zinc to a high confidence magnesium. **k**, Superimposed ion coordination site in four apoferritin structures: 7A4M (green), 7RRP (purple), 7A6A (teal), 8J5A (yellow). An additional site is shown in the top left, assigned sodium in 7RRP and water in all other structures.

On the other end of the resolution spectrum are the atomic-resolution structures of apoferritin determined by several labs,^66–69^ generally producing quite superimposable structures (**Fig. 3i**), although not without some disagreements in ion modeling. In four examples (PDB:7A4M, 7RRP, 7A6A, 8J5A) a common coordination site near glutamate 27 and 62 is modeled as either sodium (7A6A, 8J5A) or zinc (7A4M, 7RRP) (**Fig. 3j**). Interestingly in 3 of these cases (7A6A, 8J5A, 7A4M) MIC suggests a 70% or greater probability of zinc, while in 7RRP, where this site is modeled as zinc, MIC predicts a 71.2% chance of magnesium. Although the generally short coordination distances (1.9-2.1 Å) of two glutamates and a histidine support the choice of zinc in 7A4M, 7A6A, and 8J5A, the slight outward rotation and imidazole flip of histidine 65 in 7RRP weakens the case for zinc substantially as this interaction is abolished (it should be noted that in the case of 7A4M there is an alternative conformation for histidine 65 that matches 7RRP, however MIC only considers the first alternate conformation for a residue). 7RRP also includes several other ions not found in the other structures, including a zinc interacting with arginine 22 that, given the mismatched charges, should likely be a water or chloride and is predicted by MIC as water with 92.8% confidence (**Fig. 3k**). A sodium ion is also modeled interacting with the same arginine in 7RRP (**Fig. 3k**), which is similarly predicted by MIC to be a water with 97.8% confidence. These structures also have numerous waters modeled, and at this extremely high resolution, it is even possible at some sites to observe the slight deformation of the spherical densities due to the water hydrogens, providing experimental evidence for the water in some cases. Examining the 110 water molecules modeled in 7A4M, 106 (96.4%) are predicted to be water by MIC with an average confidence of 87.8±13.4%. Two sites are labeled as chloride at modest confidence (51.8% Cl, 47.3% water for A:HOH:380 and 78.9% Cl, 19.3% water for A:HOH:391, **Supplementary Table 4)**, which is possible given their interactions but there is not enough evidence for the swap. The other two discrepant sites are immediately adjacent to the zinc site, and are also assigned to be cations (sodium and magnesium) at low confidence (43.7-49.8%). This is a consistent pathology we have observed with MIC for proteins, which is that water molecules that are part of the coordination sphere of a cation are often annotated as cations with low confidence (**Extended Data Fig. 2c**). This likely stems from the fact that the model is blinded to the identity of the other nearby sites, and waters that are part of a cation coordination shell often have relatively short distances to several anionic side chains and potential ion sites themselves. To account for this, MIC warns when a site is part of a dense cluster of other sites to examine the central, high confidence site as the probable ion. Overall, the MIC method performs well for the cryo-EM structures, especially those obtained at very high resolution.

### RNA/Ribosomal structure evaluation

We wanted to examine the performance of MIC on structures of RNA, where ion binding is also pivotal,^2^ but only 72 of the 10,364 individual structures in the prevalent-ion training set contained RNA or RNA/protein complexes, corresponding to 122 ion/water sites. In general, MIC was still able to perform reasonably well on RNA-bound ions in simple high resolution RNA structures, likely correctly predicting 8/9 ions in 8D2B, 2/2 ions in 5HNJ, and 8/9 non-potassium ions in 1L2X (**Table 1**). This includes in some cases probable corrections, for example predicting the three sodium ions in 1L2X to have a strong potential to be water (**Fig. 4a-d**). This result is consistent with the overall long coordination distances for a sodium (generally 2.7-2.8 Å vs 2.4 Å expected) and the lack of more than 2 definitive hydrogen bond acceptors or 4 interaction partners total. However, where the model has more difficulties in RNA-bound structures are water molecules, which tend to be overpredicted as cations. For 1L2X, MIC had 73.8% accuracy over the 160 waters with 75.6±1.6% confidence for correct assignments and 56.5±0.13% confidence for incorrect, demonstrating both less accurate and less confident guesses, with every misassignment either sodium or magnesium. Indeed, even the sodium ions in 1L2X likely correctly predicted to be water only have ∼50% confidence.

**Fig. 4:**
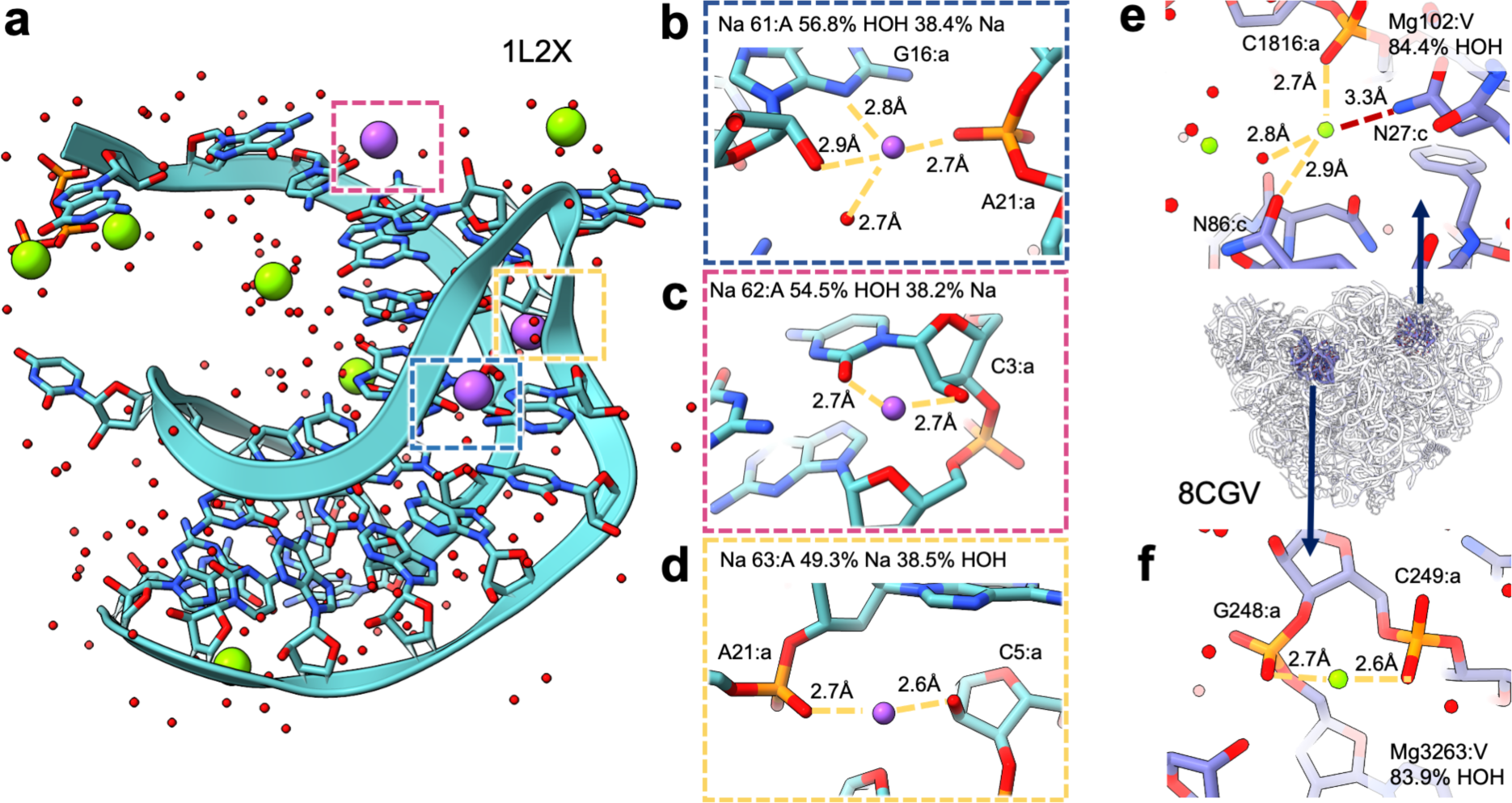
Predictions on RNA/Ribosomal structures. **a,** Structure of viral RNA pseudoknot (PDB: 1L2X). **b-d,** Sodium sites with either low-confidence water **(b,c)** or low-confidence sodium **(d)** MIC predictions. **e,f,** Potentially mismodeled magnesium ions in PDB 8CGV, predicted to be water with high confidence. **e,** MG:V:102. **f,** MG:A:3263.

**Table 1.**
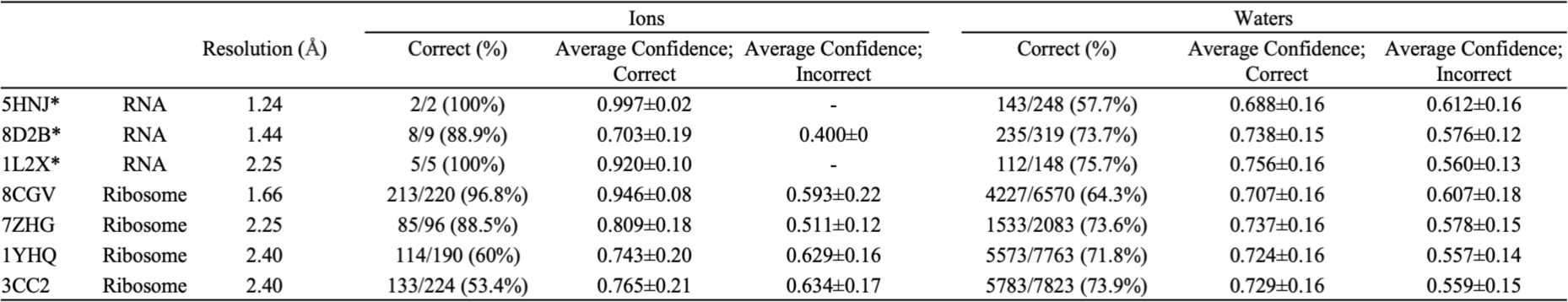
Summary of RNA/Ribosome structure performance. * indicates structures for which all non-water sites were manually examined.

This trend persists when evaluating MIC on ribosomal structures. In the case of 8CGV, the bacterial 50S ribosome at 1.66 Å resolution, MIC correctly predicts 212/219 magnesium with an average of 94.5±8.3% confidence (although some, such as MG:V:102 and MG:A:3263 which are predicted to be water, are likely mismodeled, **Fig. 4e,f**), the sole zinc correctly with 99.9% confidence, but only 4,227/6,570 water molecules with an average confidence of 70.7±16.0% (**Supplementary Table 5**). We anticipate this is likely due to the relative paucity of training data (only 56 waters in the training set are from RNA-containing structures) and will improve with further model training on additional deposited structures.

## Extended set model training, performance, and manual review

Another potential pitfall highlighted in the RNA work is the lack of inclusion of potassium or other less well-represented ions in the PDB that nevertheless can be found in structures, as the prevalent-ion model is incapable of producing the correct answer in these cases. We trained an additional model that includes potassium, iron, manganese, bromide, and iodide in addition to the prevalent ions, although there were less than 1,000 examples of each of these new classes (**Extended Data Fig. 1a, Extended Data Fig. 4a,b**). This extended-set model achieves an initial accuracy of 69.3% against the deposited test labels and displays similar results to the prevalent-ion model in embedding space organization and accuracy by class. The embedding

Similar to the prevalent set, we manually reviewed the set of discrepant ions in the extended test set using the protocol described above (**Supplementary Table 6, Extended Data Fig. 5**). This included 415 examples that were predicted to belong to a class different from the deposited label by both the prevalent and extended models as well as an additional 161 disagreeing sites belonging to the added extended classes. We observed a number of similar trends, such as a large number of sodium sites and 12 of the 86 potassium sites corrected to water in our dataset, suggesting that potassium may also be misannotated throughout the PDB (**Fig. 5e, Extended Data Fig. 5**). Even when the MIC prediction is incorrect, it can often still help identify likely changes, such as predicting the magnesium in 4AK8 to be a bromide (63.3% confidence), while the true identity is likely chloride. The final accuracy of the extended set model following manual review was 76.3%, and confidence was once again a strong measure of correctness for many classes in the prevalent set (zinc, magnesium, water, calcium) and newly introduced classes (potassium, iodide, iron, and bromine) (**Fig. 5c,d**). We observe worse chloride performance compared to the prevalent-only model, likely from the inclusion of additional halide classes that remain difficult to differentiate due to the low number of training examples. Despite this overall slight decrease in accuracy from the prevalent-only model, it is still able to successfully classify sites belonging to many different ions and can be used when one of these additional ion classes is likely.

**Fig. 5:**
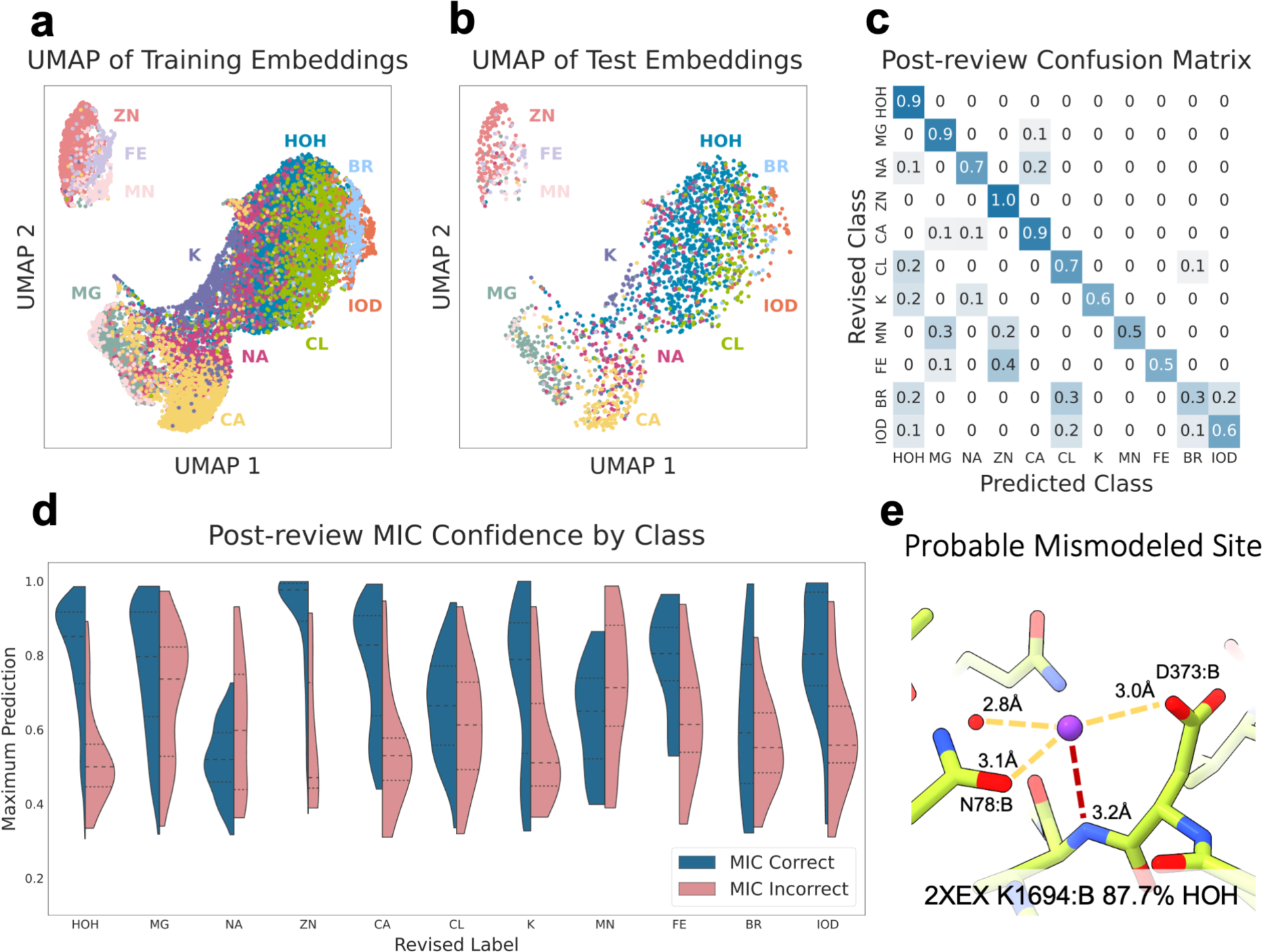
MIC Extended ion set performance and manual review. **a-b,** UMAP visualization of training **(a)** and test **(b)** set embeddings from trained extended-set MIC model. **c,** Confusion matrix of MIC predictions vs revised label following manual review. **d,** Violin plots of the confidence of correct vs incorrect MIC test set predictions by class. **e,** Probable mismodeling of a potassium site in PDB2XEX, predicted as water by MIC with 87.7% confidence. space is again organized primarily by charge as visualized by the UMAP and confirmed by PCA, transitioning smoothly from the halides to water, to monocations, and ending with the transition metals (Fig. 5a,b, Extended Data Fig 5f-h). Mis-predictions on the test set were often chemically reasonable, such as predicting bromide as either chloride or iodide, iron as zinc, or manganese as magnesium (**Extended Data Fig. 4c**). Among the added classes, iodide shows high AUROC and AUPRC values as well as separation between the confidence values of agreeing and disagreeing predictions (**Extended Data Fig. 4e,j,k**).

### Comparison with existing methods

The most ubiquitous method currently used to assign identities to ion sites is the CheckMyMetal (CMM) web server.^13–16^ CMM uses a combination of known binding site properties to evaluate each input structure. Each property (atomic contacts, valence, and geometry) contributes a score between 0 and 2 resulting in a maximum score of 6 for a given ion identity at a particular site. The score of each potential metal is reported, often leading to multiple ions receiving comparably high scores. During manual inspection of the disagreeing test examples, we ran CMM on all structures to identify cases where CMM and MIC differ in either their predicted class or from the deposited label in the PDB.

CMM and MIC produce concordant results for many sites. In 3FOB, MIC predicts the deposited sodium ion to be a magnesium (88.6%), consistent with the respective scores of 4 and 6 for these metals from CMM, although cobalt and manganese also receive a CMM score of 6 (**Fig. 6a**). Similarly, both MIC and CMM predict the calcium site in 1XPH to be a magnesium, receiving a CMM score of 6 and a MIC confidence of 82.4%. In sites where manual inspection revealed the deposited metal was likely water or chloride, CMM generally gave poor scores for either all metals or all metals except potassium. The sites of magnesium to chloride corrections in 4KP1, 3S70, and 3A4X, and the magnesium to water correction in 5VX0, receive a 0 from CMM for all ions, but no additional distinction between these cases is provided as CMM is specifically designed for validating metals. Additionally, there are cases where MIC likely identifies the correct label while CMM does not, notably in correcting ions to water: 6RJ4 is one such example containing a deposited sodium ion. CMM gives this site a potassium score of 6, followed by a sodium score of 4, showing significant disagreement with the high-confidence MIC water prediction (95.8%) that was accepted upon manual review, as re-refinement with water at this position improves both the difference map and interactions at this site (**Fig. 6b**). Conversely, because CMM predicts the output for multiple classes, it can in some cases succeed where MIC fails; the identity of the magnesium ion in 2ICJ was confirmed experimentally^70^ and agrees with the high CMM assignment of 5 for magnesium, while MIC predicts this site to contain a high-confidence zinc ion, which only scored a 4 by CMM. Finally, CMM and MIC have distinct output classes that make each more suitable for specific cases. CMM predicts the score for several metals that are not in the MIC class set, including copper, cobalt, and nickel, while MIC includes an explicit prediction for water and halides.

**Fig. 6:**
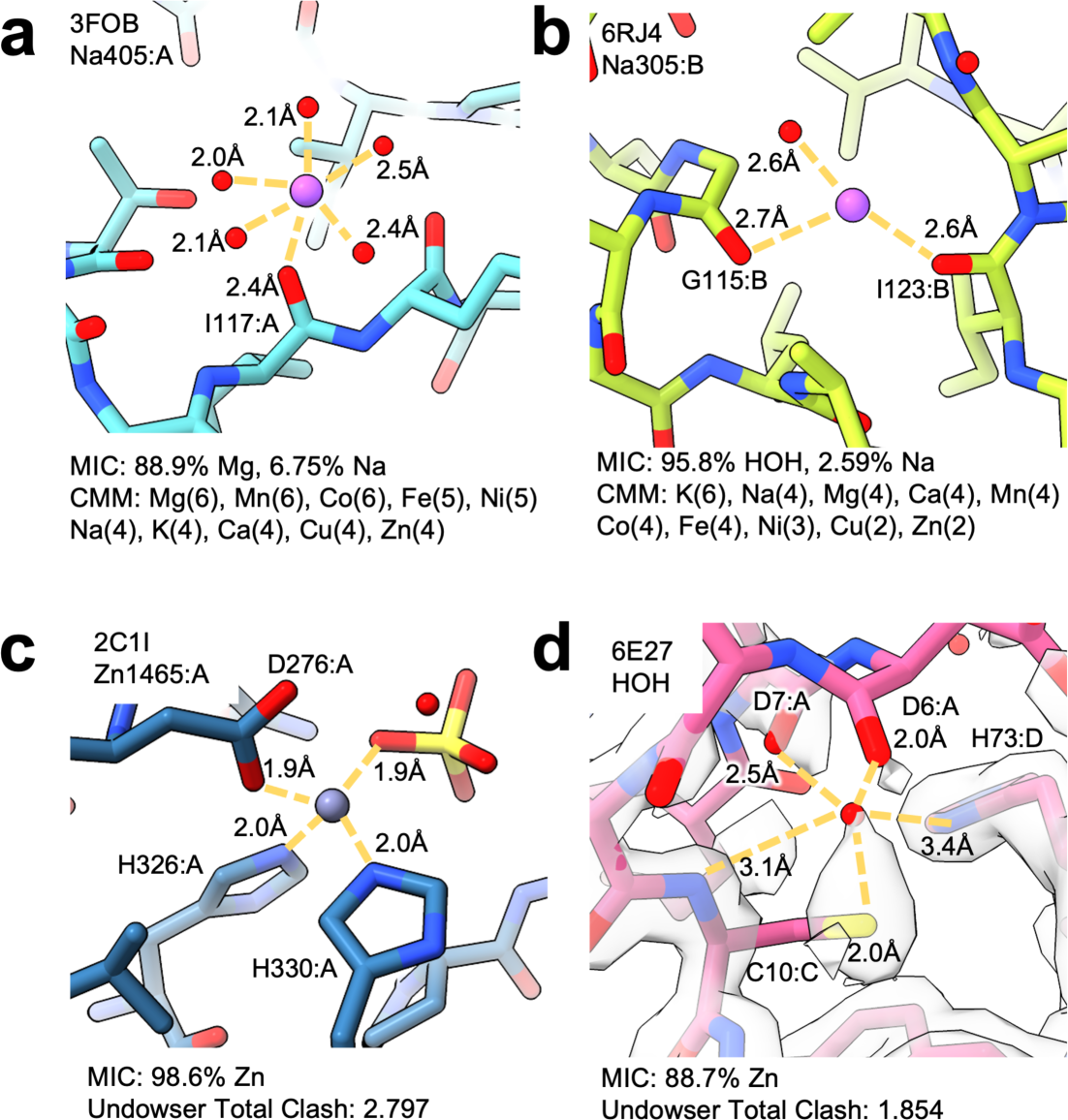
MIC, CheckMyMetal, and Undowser performance examples. **a,** Example of CMM and MIC both correcting a sodium (3FOB:A:Na:405) to a magnesium (MIC confidence: 88.9%, CMM Mg Score: 6). **b,** Example of a site (6RJ4:A:Na:305) MIC likely predicts correctly as a water (95.8% confidence) over CMM, which assigns a score of 6 for potassium and 4 for five other metals. **c**, Zinc coordination site (2C1I:A:Zn:1465) identified by both MIC and Undowser with high confidence (MIC zinc confidence: 98.6%, Undowser clash score: 2.797). **d,** Example of Undowser and MIC results at a questionably modeled site (6E27:C:HOH:201). This site likely does not contain either an ion or a water, but is predicted to be an ion by both Undowser (Clash score: 1.854) and MIC (zinc confidence: 88.7%).

Another tool with some overlapping use is Undowser, which is intended to find waters that clash with nearby atoms as these could indicate that the site would better be modeled as a metal. We ran MIC and Undowser on a selection of identity-blinded waters and ions to compare the results (see Methods, **Supplementary Table 7**). Like CMM, Undowser often agrees with MIC predictions. Both tools identify the zinc sites in 2C1I (A:1465,1466) with a MIC zinc confidence of 98.6% and 91.4%, and an Undowser cumulative clash severity score of 2.797 and 2.121, respectively, each comprising of multiple >0.5Å polar clashes strongly indicating the presence of an ion (**Fig. 6c**). Similarly, the magnesium ion in 4RKQ is caught by both tools (MIC: 98.1%, Undowser: 2.098). Even when MIC is unable to predict the correct identity, it is often able to distinguish what should be an ion binding site, such as the iron sites in 1YFU, predicted as zinc by the MIC prevalent-only model with 98.6% confidence and an Undowser clash severity score of 1.985. MIC does show a tendency to over-predict ions compared to Undowser, though similar to the RNA/ribosomal predictions these assignments typically have a lower confidence (57.1±15.1%) than water predictions that agree with Undowser (85.5±13%), helping the user handle these cases. Undowser and MIC both fail where the modeling is questionable, as is the case for 6E27:C:HOH:201, which both MIC and Undowser flag with high zinc confidence and clash score (88.7%, 1.854Å). However, 6E27 shows no major positive difference density and lacks density in the 2Fo-Fc map for much of the protein at this site (**Fig. 6d**). Undowser provides information about the charge of clashing atoms that can assist the user in interpreting the results, but does not explicitly attempt to predict the true ion identity. Undowser does not calculate any clashes for halides such as the chlorines in 3MUJ and 4RKQ, which are predicted correctly by MIC. Ultimately, these are complementary but not overlapping methods of confirming correct modeling, and users should choose the tool that best aligns with their specific requirements.

## Discussion

MIC is a novel method to classify water/ion sites in biomolecular structures and provide confidence estimation in these predictions. It uses structured embeddings from a deep metric learning model to generate accurate predictions for structures from both x-ray crystallography and cryo-EM. Notably, this representation includes no angle information and only coarse distance information through constructing the initial microenvironment fingerprint by shell expansion. Despite this, our results show that the fingerprints contain sufficient data about the atomic environment to correctly distinguish ions in over 80% of our observed cases. We demonstrate that MIC achieves incredible accuracy for extremely high-resolution cryo-EM structures. Furthermore, the results for MC4R suggest that ambiguity and inaccurate assignments by MIC may result from problematic protein modeling. The output probabilities can be used to estimate model confidence, further enabling MIC’s use as a validation tool. We use MIC to identify mismodeled ions throughout the PDB and show that the tool can be used to evaluate the modeling of coordinating side-chains in low-resolution structures when the desired ion identity is known. Finally, we show that MIC performs comparably to field-standard approaches and offers further utility over these methods by including additional output classes.

There are limitations to MIC’s use. Our dataset was restricted to ions for which at least a few hundred high-quality structures exist in the PDB, limiting the possible classes we could include when training the deep-learning model and SVC portions. As mentioned previously, there are concerns about the quality of the training dataset, specifically the inclusion of misannotated ions from the PDB. Sodium in particular achieves lower performance and even correct sodium predictions are often low confidence, likely due to mislabeled sodium sites introducing noise into the dataset. MIC also tends to predict waters that are very close to a cation and participate in a larger coordination cluster as cations, necessitating an additional flag to account for these cases. MIC is designed to work for sites centered in experimentally determined density, and is not expected to give accurate results for sites with few or no interactions.

The current MIC workflow uses a deterministic hash function to convert the site microenvironment to a fixed-length fingerprint as input to the deep-learning model. This introduces the potential for bit collisions that occur when multiple features hash to the same index in the fingerprint, complicating both model training and feature attribution analyses. A future expansion on this method could replace this fingerprinting method with a task-specific representation learned from this proximity graph, similar to those used for molecular property prediction,^71–73^ further enabling downstream classification and limiting the problem of bit collisions. The metric model was trained using a triplet loss to maximize inter-class distances and successfully learns class relationships, but this could be explicitly enforced using a hierarchical loss function. As new structures are added to the Protein Data Bank and ion sites are subject to more careful validation and scrutiny, this additional training data will further improve MIC’s accuracy for future iterations.

## Methods

### Dataset Curation

Our final dataset consisted of 23,101 examples split across 11 classes. In decreasing order of number of examples: water, magnesium, sodium, zinc, calcium, chloride, potassium, manganese, iodine, iron, and bromine. Candidate structures were restricted to < 2.0 Å resolution except for potassium, bromide, and iodide, which were relaxed to < 3.0 Å. Structures were restricted to have no more than 95% sequence homology to another example in the dataset to avoid overrepresentation. Only the first example of each ion type and a single water was taken from a given structure to prevent redundant symmetry-related ions being added to the dataset and potential modeler-related biases. Finally, to prevent the inclusion of water molecules and/or ions modeled into empty space with no interaction partners, sites were filtered to have at least 2 other atoms within 3.5 Å. The full list of ions and their associated counts and resolutions is provided in **Extended Data Fig. 1a-b** and all sites used along with the training and testing splits are provided in **Supplementary Table 1**.

### Density Fingerprint Representation

We represented each density using a modified version of the interaction fingerprint available in the LUNA toolkit. LUNA fingerprints were developed to capture the interactions at the protein-ligand interface of a bound complex. This is accomplished by assigning biochemical properties of individual atoms and atomic groups, then defining interactions as pairs of atoms/atomic groups in proximity that meet certain geometric and chemical properties. The ligand properties and final list of intermolecular interactions are then converted into a fixed length vector using the MurmurHash3 algorithm, referred to as a fingerprint.^35,74^

We made two crucial modifications to the generation protocol of these interface fingerprints for our use case. First, to ensure that any previous density assignment was not encoded in the representation, trivializing any downstream classification task, the initial atomic representation for each density of interest and nearby spherical densities was programmatically set to a vector of zeros. All atoms belonging to the protein or nearby small molecules are given their standard initial feature set (described in more detail below). The second modification was limiting all calculated interactions to be proximal-only, defined as simply being located between 2Å and 6Å from the ion. Proximal interactions are unique in that they do not rely on the chemical features of either participating atomic group, and thus the resulting representation continues to be identity-agnostic. The final fingerprint is effectively an identity-blinded representation of the atomic environment surrounding the density of interest.

### Initial atomic features and included interactions

LUNA provides extended interaction fingerprint (EIFP) and functional interaction fingerprint (FIFP) featurization options for the user during fingerprint generation. EIFPs use the Daylight atomic invariants for the initial atomic feature set, consisting of 7 fields: number of heavy atom neighbors, valence minus the number of bound hydrogens, atomic number, isotope number, formal charge, number of hydrogen neighbors, and aromaticity.^75^ Functional fingerprints use pharmacophore-like features, such as whether an atom is aromatic, hydrophobic, or a hydrogen donor or acceptor. **Extended Data Table 1** contains the full list of initial atomic features for each fingerprint type. In addition to these options, we also considered whether or not to include interactions between neighbors in our representation versus a “pruned” representation with interactions limited only to those between the density of interest and neighbors. We evaluated the four fingerprint types (non-prune/eifp, prune/eifp, non/prune-fifp, prune/fifp) by generating the specified fingerprint type and training an SVC to predict ion identity. We found that prune-eifp fingerprints performance exceeded that of the other types, especially in multi-class classification tasks, and used that type for all presented work (**Extended Data Fig. 1c-f)**.

### Shell Number and Radius

During the fingerprint creation process, interactions in shells around the ligand atoms are iteratively converted to fixed integers, similar to the process for generating molecular Morgan fingerprints.^35^ The user sets the size and number of these shells during creation, with the default LUNA values being 2 shells of 6Å radius step each. We hypothesized that these values, optimized for longer intermolecular protein-ligand interfaces, would not provide sufficient granularity to differentiate between ions. To address this, we explored a range of different shell radii and depths by randomly selecting up to 2000 examples from each ion class in the dataset, generating fingerprints with the specific radii and number of shells, and training an SVC^76^ with 5-fold cross-validation to predict ion identity (**Extended Data Fig. 1c-f**). Accuracies ranged from 0.44 to 0.63 for all fingerprint types, with a general rule that the product of the radius and number of shells should be between 3Å and 5Å for best performance. Lower radii performed better overall, suggesting that the additional discrimination provided by finer shells is useful for downstream identity classification, though this does make the representation more sensitive to slight changes in atom position. All final fingerprints were generated with 18 shells of radius step 0.25Å, comprising a total volume of 4.5Å around each atom in the proximity graph. Count fingerprints of length 4,096 were used for all experiments, consistent with the original LUNA manuscript.

### Training and test datasets of curated densities for MIC

Each site in our curated dataset was randomly assigned to training or testing with a 90%/10% split. We chose random splits because of the strict criteria during dataset curation, limiting the similarity between all examples. All ions belonging to a class with fewer than 1000 examples were dropped for the prevalent-only models, resulting in final datasets of 18,420 training and 2037 testing examples. The extended set included all of these sites plus examples from potassium, manganese, iron, bromide, and iodide, bringing the total number of training examples to 20,801 with 2,300 examples used for testing. Training and testing splits were consistent across all fingerprint types and hyperparameter optimization.

### Model Training

We present two metric learning networks one trained on the prevalent set of ions and one trained on the extended set. Models were trained using the Pytorch Metric Learning library with triplet margin sampling and triplet loss.^77,78^ Hyperparameter optimization was performed with the Optuna library to evaluate the effect of learning rate, dropout, loss and miner margins, and embedding dimension.^79^ The full list of evaluated hyperparameters, ranges, and final values are displayed in **Extended Data Table 2**. Models were optimized to maximize two downstream metrics: the average area under the receiver operating characteristic (ROC) curve and the F1-score for a linear-kernel SVC trained on the embeddings from the metric learning model. Due to significant class imbalance, each batch was generated by weighted random sampling with replacement, resulting in approximately balanced batches. The architecture of all final models consists of 1 hidden layer of 4000 neurons each with output size of 32 for the resulting embeddings. The model was trained for 1000 epochs (**Extended Data Fig. 2a,b**).

### Feature Attribution

Pairwise feature attribution was calculated between representative examples for each class using a modification of the popular integrated gradients technique implemented in the Captum library.^57,58^ The specific examples were chosen by selecting those close to the cluster centroid for a given class in the learned embedding space. A baseline fingerprint of zeroes was used for all calculations. Following default global attribution rules, the resulting attribution vector is multiplied by the input fingerprint. While this is known in practice to result in cleaner attribution features and improve the ease of interpretation of the results, it is an important limitation to note as only features that are turned “on” for a given fingerprint will be assigned a non-zero attribution value. The features corresponding to the top ten bits with the highest attribution for both comparisons are available in **Supplementary Table 3.**

### Undowser Comparison

The comparison with Undowser was performed by randomly selecting structures from the PDB that fit our resolution requirements and contained at least one non-water density, converting all of the ions to water, and running Undowser to determine if the clashing “waters” matched with the non-water MIC predictions.

### Statistical Analysis

All statistical analyses performed were two-sided t-tests for independence, as implemented in the Scipy^80^ statistics module. The full list of comparisons, number of examples, and P-values are provided in **Extended Data Table 3**.

## Data and code availability

The complete source code for MIC, all training data, trained models, and associated tutorial Jupyter Notebooks are freely available under the open-source MIT license at https://github.com/keiserlab/metric-ion-classification.

## Acknowledgements

This work was supported by NIH T32 GM067547 and the UCSF Graduate Division (L.S), CZI grant DAF2018-191905 (DOI 10.37921/550142lkcjzw) from the Chan Zuckerberg Initiative DAF, an advised fund of Silicon Valley Community Foundation (funder DOI 10.13039/100014989) (M.J.K.), NIH K99/R00 HD107581 (M.J.R.), and CPRIT award RR230042 (M.J.R.). We would like to thank Alexandre Fassio for his assistance in adapting the LUNA package, Luca Ponzoni for his work and advice in implementing feature attribution, and Mahdi Ghorbani, Brendan Hall, and Zachary Gale-Day for their useful advice throughout this project.

## Author contributions

L.S. designed and wrote the MIC software, trained and evaluated the machine learning models, and generated predictions for all structures. M.J.R. conceived the project, curated training data, and performed manual validation and structure re-refinement. S.L. provided code review and software validation. L.S. and M.J.R. wrote the manuscript with input from G.S., M.J.K., and S.L. M.J.R, G.S., and M.J.K. supervised the project.

---

**Extended Data Figure 1.**
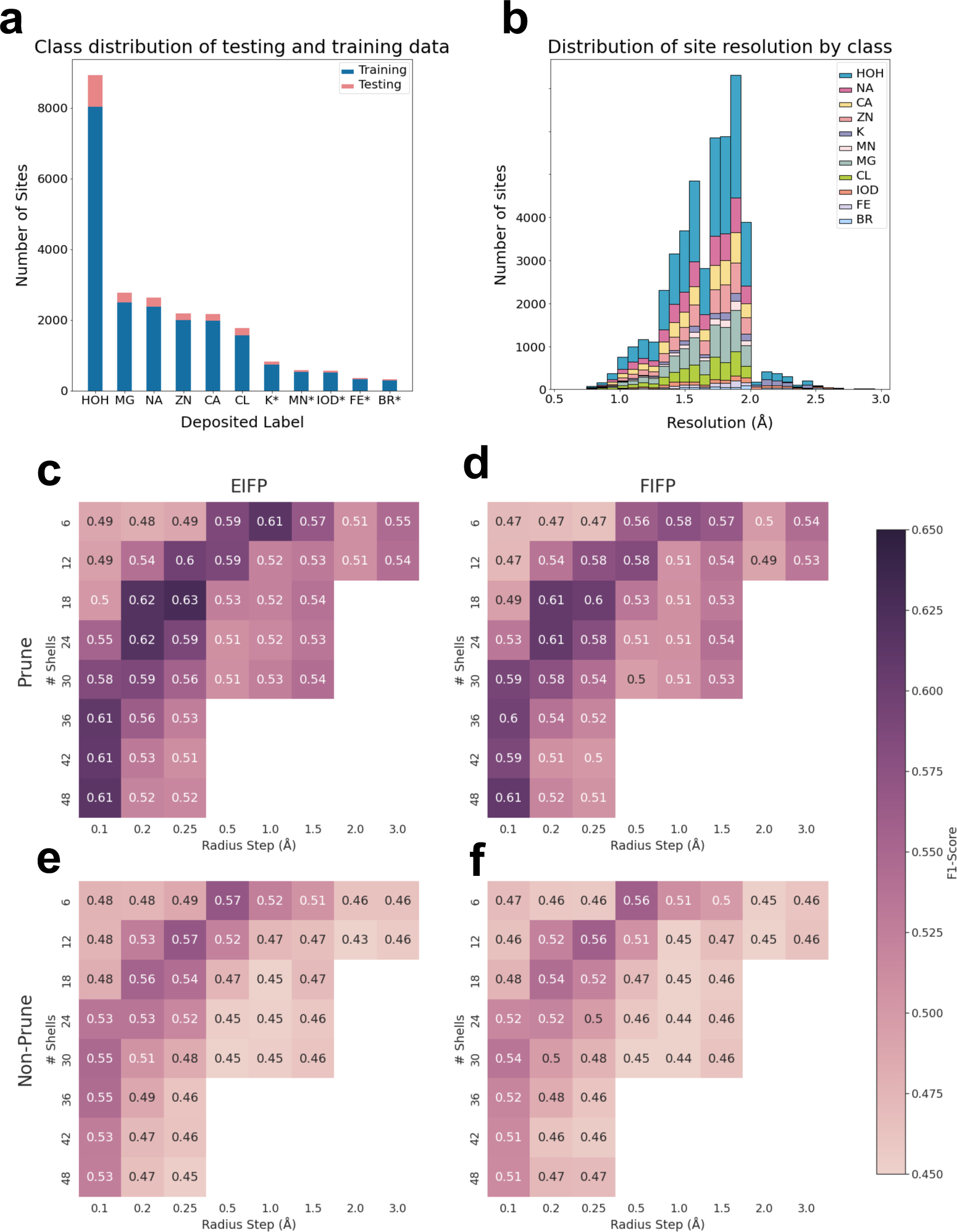
MIC dataset preparation and exploration. **(a)** Class distribution used for training and testing MIC models. Extended-set only ions are marked with *. **(b)** Distribution of the resolutions of the structures each site was extracted from, colored by the deposited label. Potassium, chloride, and bromide (and matching waters) were restricted to <3Å, all other classes were restricted to <2Å. **(c-f)** Heatmaps of the accuracy of an SVC trained directly on fingerprints generated with a given radius and shell number for the evaluated fingerprint types. **(c)** Prune/ EIFP, **(d)** Non-prune /EIFP, **(e)** Prune/FIFP, **(f)** Non-prune/FIFP

**Extended Data Figure 2.**
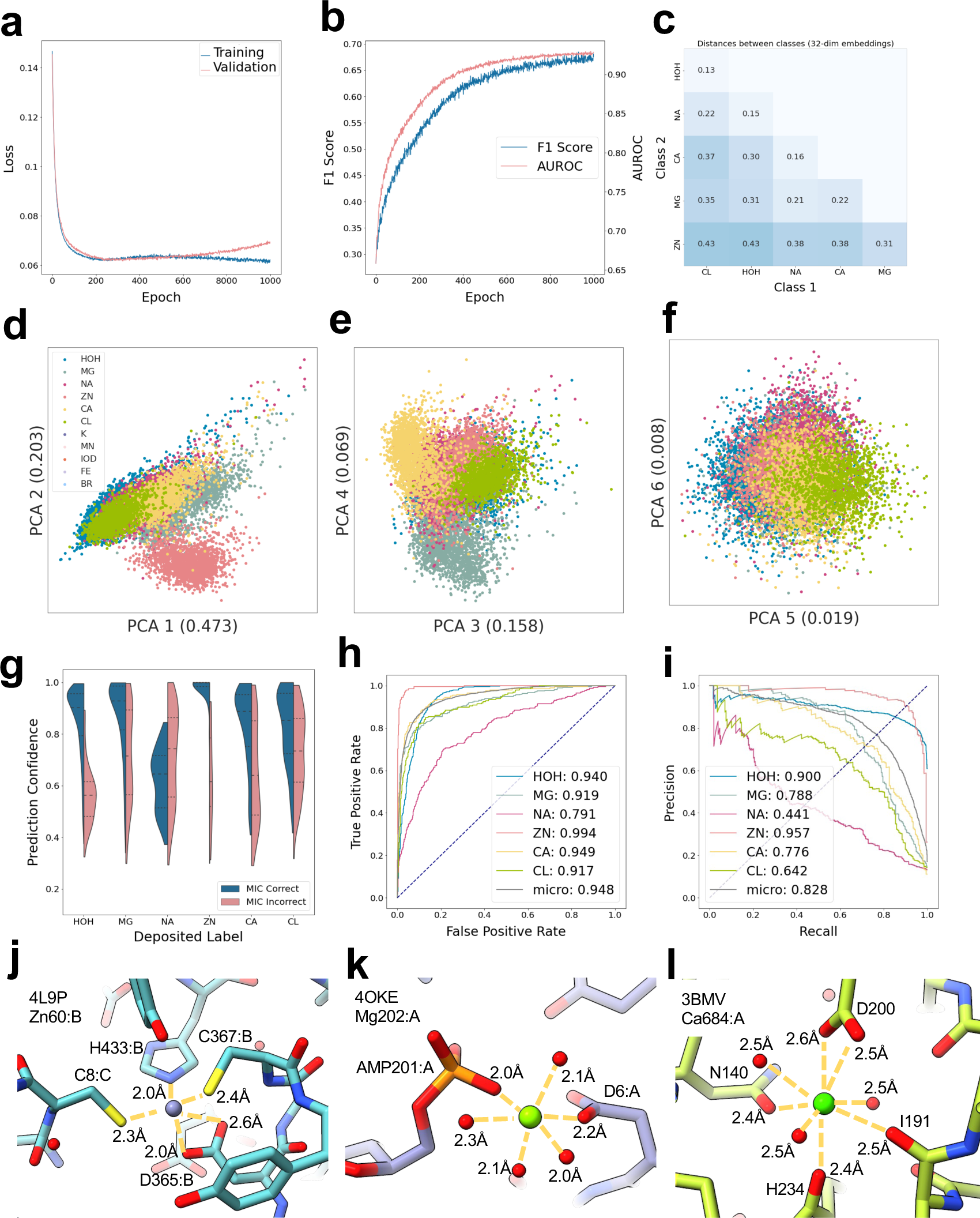
MIC prevalent-ion set additional results. **(a)** Training and validation loss. **(b)** F1-score and micro-AUROC of SVC trained on intermediate training set embeddings and labels, evaluated on validation set. **(c)** Average distance in 32-dim latent embedding spaces between classes as confirmation of the trends observed in the Fig. 2 Low-dimensional visualizations. **(d-f)** PCA plots for of the first 6 dimensions by variance explained, shown alongside the axis labels. **(g)** Confidence of agreeing vs disagreeing predictions and deposited labels of prevalent-ion test set, split by class. **(h)** ROC curves of individual classes and micro-average of MIC predictions on the prevalent-ion test set. **(i)** PRCs of individual classes and micro-average of MIC predictions on the prevalent-ion test set. **(j-l)** Sites used to perform feature attribution analysis. **(j)** 4L9P:B:ZN:601. **(k)** 4OKE:A:MG:202. **(l)** 3BMV:CA:A:684.

**Extended Data Figure 3.**
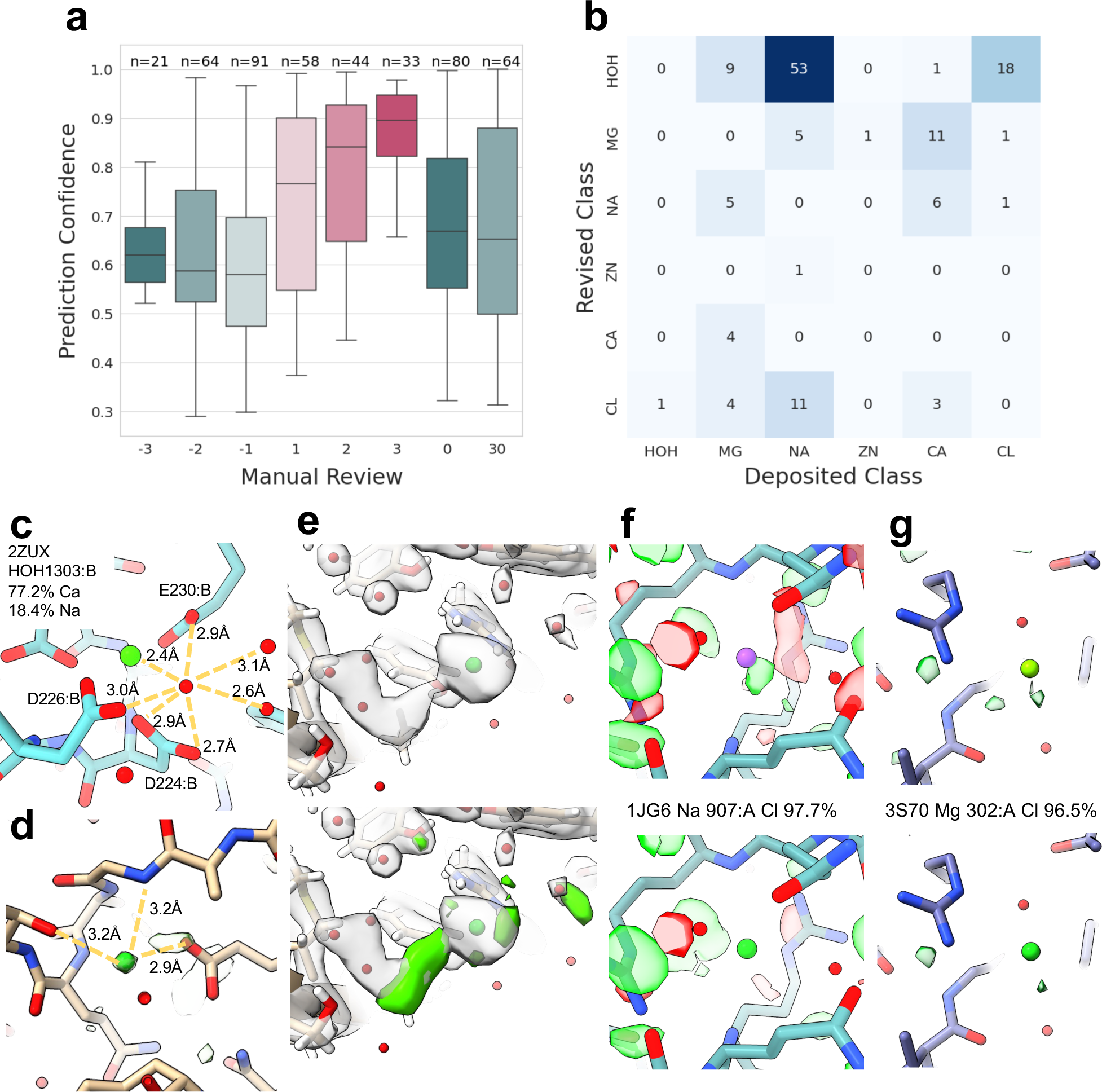
Manual review of test set discrepant sites. **(a)** Confidence of each discrepant example vs the manual review value. Manual review values ranged from -3, indicating an incorrect MIC prediction to 3, indicating a correct MIC prediction. 0 marks examples for which it was unclear which of the two labels was correct, and 30 marks sites with other issues preventing proper prediction. **(b)** Heatmap of corrected sites; deposited label is shown on the x-axis and the revised MIC-predicted label is shown on the y-axis. Each cell is labeled with the count of sites. **(c)** Example of common MIC pathology of predicting coordinating waters as cations. **(d-e)** Examples of structures that received a 0 and a 30 during manual review. **(d)** 4DWD. **(e)** 4JO5. **(f-g)** Re-refinement of difference maps contoured at +/- 3σ with deposited (top) and MIC-predicted label (bottom) **(f)** 1J68. **(g)** 3S70.

**Extended Data Figure 4.**
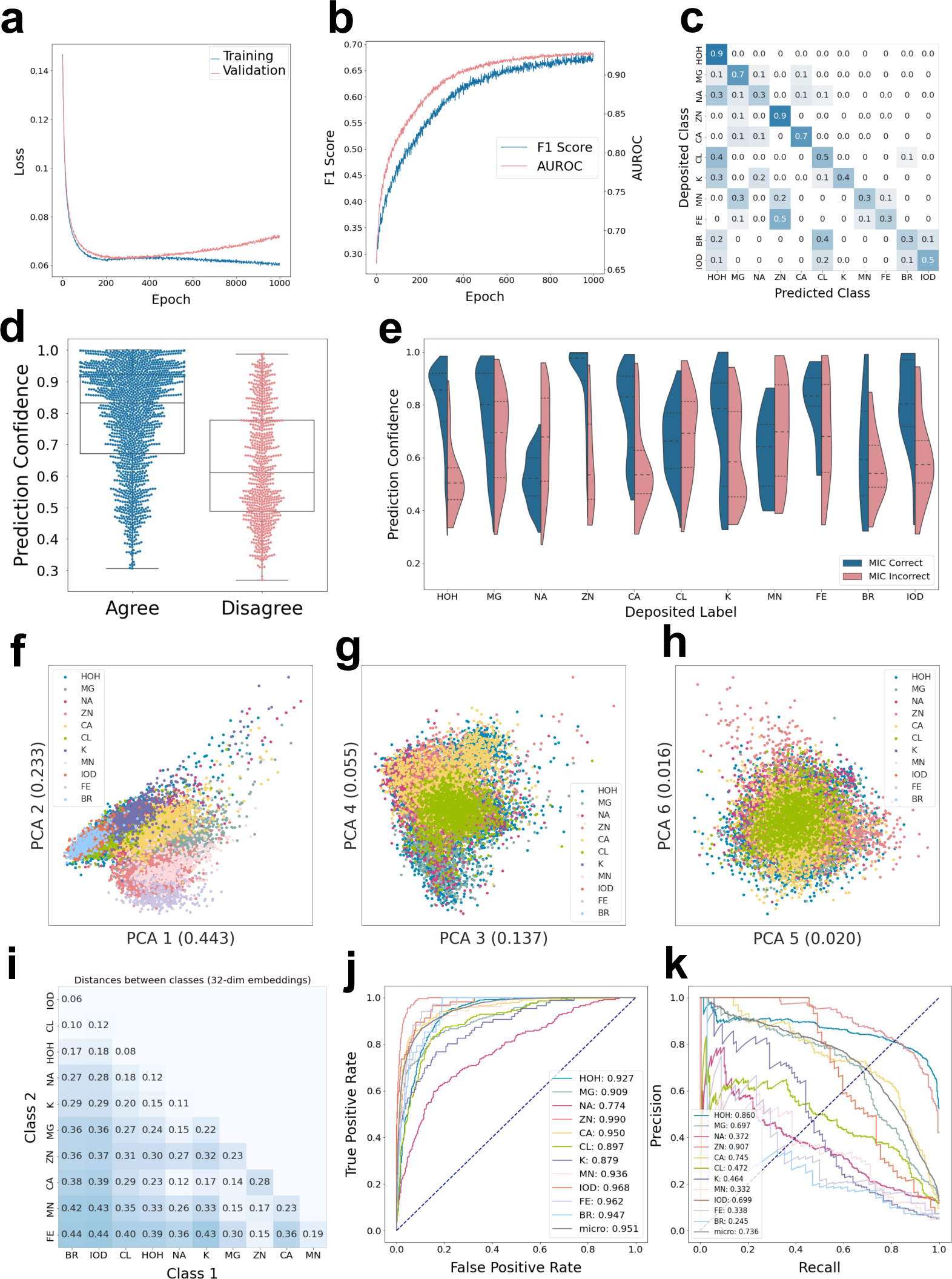
MIC extended set additional results. **(a)** Training and validation loss. **(b)** AUROC and F1-score during training. **(c)** Confusion matrix of test set deposited and predicted label. **(d)** Agreeing vs discrepant prediction confidence, pre-revision. **(e)** Agreeing vs discrepant prediction confidence by class, pre-revision. **(f-h)** PCA plots of learned latent embeddings, extended set. **(i)** Distance between classes in embedding space. **(j)** ROC curves of individual classes and micro-average of MIC predictions, extended set. **(k)** PRCs of individual classes and micro-average of MIC predictions, extended set.

**Extended Data Figure 5.**
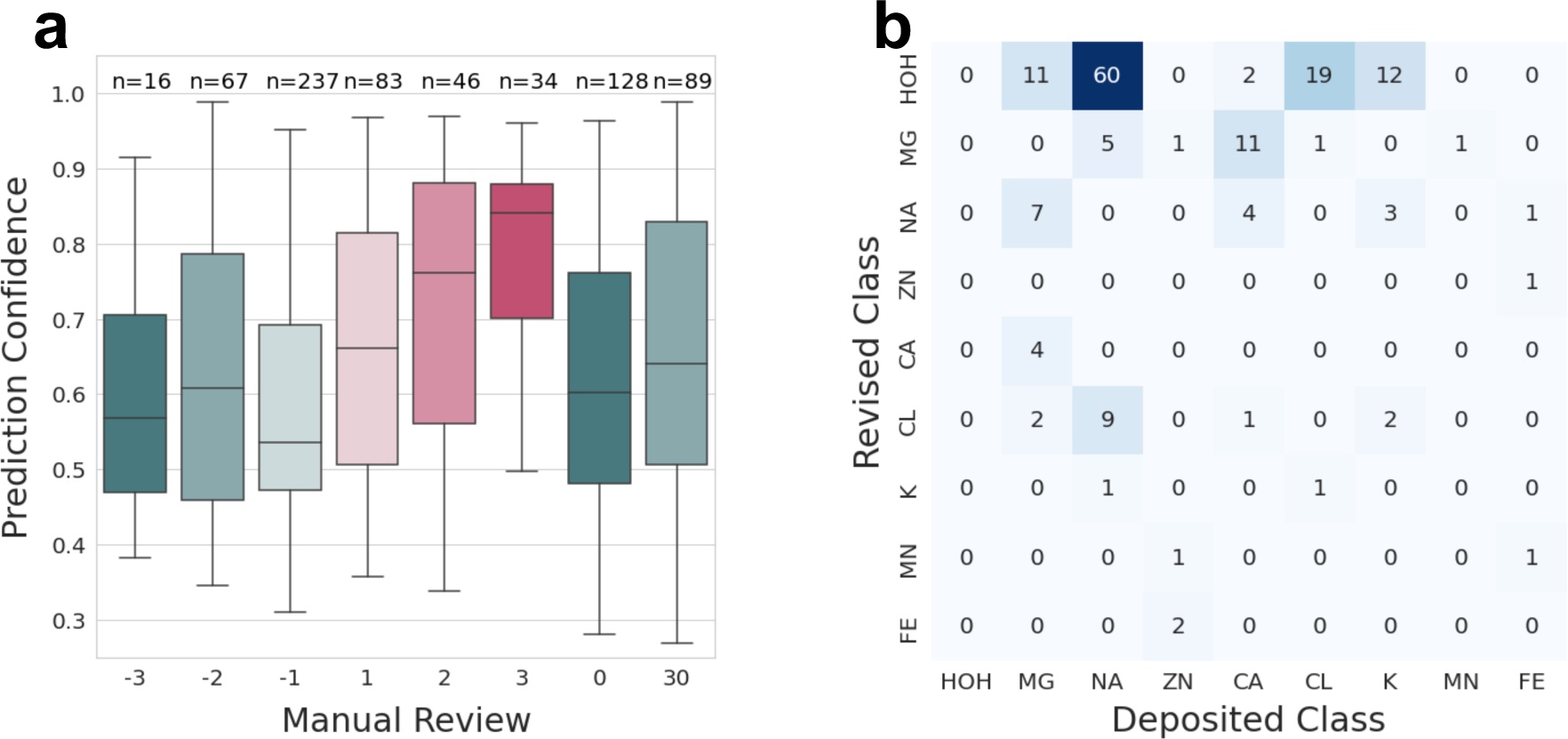
Manual review of discrepant sites from the extended test set. **(a)** Confidence of each discrepant example vs the manual review value. Manual review values ranged from -3, indicating an incorrect MIC prediction to 3, indicating a correct MIC prediction. 0 marks examples for which it was unclear which of the two labels was correct, and 30 marks sites with other issues preventing proper prediction. **(b)** Heatmap of corrected sites; deposited label is shown on the x-axis and the revised MIC-predicted label is shown on the y-axis. Each cell is labeled with the count of sites.

**Extended Data Table 1.** Initial features for each atom by type and fingerprint type.

| Type | Descriptors | Length |
| --- | --- | --- |
| Ions and waters | [0, 0, 0, 0, 0, 0, 0] | 7 |
| Extended Interaction Fingerprint (EIFP) | [# Heavy Atom Neighbors, Valence - # Hydrogen Neighbors, Atomic number, Isotope number, Formal Charge, # Hydrogen Neighbors, Is_In_Ring] | 7 |
| Functional Interaction Fingerprint (FIFP) | Aromatic, Acceptor, Donor, Hydrophobe, Hydrophobic, Negative, Positive, Negatively ionizable, Positively ionizable, Halogen donor, Metal, Lumped hydrophobe, Weak donor, Weak acceptor, Electrophile, Nucleophile, Chalcogen donor, Amide, Atom | $\leq 19$ |

**Extended Data Table 2.** Hyperparameters explored during optimization and values used for the final MIC models.

| Hyperparameter | Options | Final |
| --- | --- | --- |
| hidden_layers | [1,4] | 1 |
| n_neurons | 1000,2000,3000,4000 | 4000 |
| embedding_dim | 2, 8, 16, 32 | 32 |
| learning_rate | [1e-8, 1e-4] | 1e-7 |
| dropout | [0.05, 0.4] | 0.35 |
| weight_decay | [0, 0.1] | 0.005 |
| loss_margin | [0.01, 0.1] | 0.06 |
| miner_margin | [0.1, 0.3] | 0.2 |

**Extended Data Table 3.** P-values and number of sites for all statistical analysis. Statistically significant comparisons are indicated.

| Comparison | Number of examples | P-value |
| --- | --- | --- |
| Confidence of agreeing vs disagreeing predictions, initial* | 2037 | 4.802e-83 |
| Confidence of agreeing vs disagreeing predictions, revised* | 1893 | 8.596e-74 |
| Confidence of agreeing vs disagreeing water predictions, revised* | 975 | 8.166e-49 |
| Confidence of agreeing vs disagreeing magnesium predictions, revised* | 245 | 3.450e-10 |
| Confidence of agreeing vs disagreeing sodium predictions, revised | 151 | 0.178 |
| Confidence of agreeing vs disagreeing zinc predictions, revised* | 193 | 5.952e-17 |
| Confidence of agreeing vs disagreeing calcium predictions, revised* | 165 | 1.536e-06 |
| Confidence of agreeing vs disagreeing chlorine predictions, revised* | 164 | 3.197e-07 |

## References

1. Bellissent-Funel, M.-C. et al. Water Determines the Structure and Dynamics of Proteins. Chem. Rev. 116, 7673–7697 (2016).

2. Draper, D. E. A guide to ions and RNA structure. RNA 10, 335–343 (2004).

3. Lu, Y., Wang, R., Yang, C.-Y. & Wang, S. Analysis of Ligand-Bound Water Molecules in High-Resolution Crystal Structures of Protein−Ligand Complexes. J. Chem. Inf. Model. 47, 668–675 (2007).

4. Anderson, A. C. The Process of Structure-Based Drug Design. Chem. Biol. 10, 787–797 (2003).

5. Mobley, D. L. & Dill, K. A. Binding of Small-Molecule Ligands to Proteins: “What You See” Is Not Always “What You Get”. Structure 17, 489–498 (2009).

6. Andreini, C., Bertini, I., Cavallaro, G., Holliday, G. L. & Thornton, J. M. Metal ions in biological catalysis: from enzyme databases to general principles. J. Biol. Inorg. Chem. JBIC Publ. Soc. Biol. Inorg. Chem. 13, 1205–1218 (2008).

7. Hediger, M. A., Kanai, Y., You, G. & Nussberger, S. Mammalian ion-coupled solute transporters. J. Physiol. 482, 7–17 (1995).

8. Liebschner, D. et al. Polder maps: improving OMIT maps by excluding bulk solvent. Acta Crystallogr. Sect. Struct. Biol. 73, 148–157 (2017).

9. Yamashita, K., Palmer, C. M., Burnley, T. & Murshudov, G. N. Cryo-EM single-particle structure refinement and map calculation using Servalcat. Acta Crystallogr. Sect. Struct. Biol. 77, 1282–1291 (2021).

10. Wang, J., Liu, Z., Frank, J. & Moore, P. B. Identification of ions in experimental electrostatic potential maps. IUCrJ 5, 375–381 (2018).

11. Yamashita, M. M., Wesson, L., Eisenman, G. & Eisenberg, D. Where metal ions bind in proteins. Proc. Natl. Acad. Sci. 87, 5648–5652 (1990).

12. Carugo, O. Buried chloride stereochemistry in the Protein Data Bank. BMC Struct. Biol. 14, 19 (2014).

13. Gucwa, M. et al. CMM—An enhanced platform for interactive validation of metal binding sites. Protein Sci. Publ. Protein Soc. 32, e4525 (2023).

14. Handing, K. B. et al. Characterizing metal binding sites in proteins with X-ray crystallography. Nat. Protoc. 13, 1062–1090 (2018).

15. Zheng, H. et al. CheckMyMetal: a macromolecular metal-binding validation tool. Acta Crystallogr. Sect. Struct. Biol. 73, 223–233 (2017).

16. Zheng, H. et al. Validation of metal-binding sites in macromolecular structures with the CheckMyMetal web server. Nat. Protoc. 9, 156–170 (2014).

17. Prisant, M. G., Williams, C. J., Chen, V. B., Richardson, J. S. & Richardson, D. C. New tools in MolProbity validation: CaBLAM for CryoEM backbone, UnDowser to rethink “waters,” and NGL Viewer to recapture online 3D graphics. Protein Sci. 29, 315–329 (2020).

18. Jumper, J. et al. Highly accurate protein structure prediction with AlphaFold. Nature 596, 583–589 (2021).

19. Baek, M. et al. Accurate prediction of protein structures and interactions using a three-track neural network. Science 373, 871–876 (2021).

20. Watson, J. L. et al. De novo design of protein structure and function with RFdiffusion. Nature 620, 1089–1100 (2023).

21. Corso, G., Stärk, H., Jing, B., Barzilay, R. & Jaakkola, T. DiffDock: Diffusion Steps, Twists, and Turns for Molecular Docking. Preprint at 10.48550/arXiv.2210.01776 (2023).

22. Stärk, H., Ganea, O.-E., Pattanaik, L., Barzilay, R. & Jaakkola, T. EquiBind: Geometric Deep Learning for Drug Binding Structure Prediction. Preprint at 10.48550/arXiv.2202.05146 (2022).

23. Doerr, S. et al. TorchMD: A Deep Learning Framework for Molecular Simulations. J. Chem. Theory Comput. 17, 2355–2363 (2021).

24. Si, D. et al. Deep Learning to Predict Protein Backbone Structure from High-Resolution Cryo-EM Density Maps. Sci. Rep. 10, 4282 (2020).

25. Maddhuri Venkata Subramaniya, S. R., Terashi, G. & Kihara, D. Protein secondary structure detection in intermediate-resolution cryo-EM maps using deep learning. Nat. Methods 16, 911–917 (2019).

26. Terashi, G., Wang, X., Prasad, D., Nakamura, T. & Kihara, D. DeepMainmast: integrated protocol of protein structure modeling for cryo-EM with deep learning and structure prediction. Nat. Methods 21, 122–131 (2024).

27. Jamali, K. et al. Automated model building and protein identification in cryo-EM maps. Nature 1–2 (2024) doi:10.1038/s41586-024-07215-4.

28. Wang, X., Terashi, G. & Kihara, D. CryoREAD: de novo structure modeling for nucleic acids in cryo-EM maps using deep learning. Nat. Methods 20, 1739–1747 (2023).

29. Terashi, G., Wang, X., Maddhuri Venkata Subramaniya, S. R., Tesmer, J. J. G. & Kihara, D. Residue-wise local quality estimation for protein models from cryo-EM maps. Nat. Methods 19, 1116–1125 (2022).

30. He, J., Li, T. & Huang, S.-Y. Improvement of cryo-EM maps by simultaneous local and non-local deep learning. Nat. Commun. 14, 3217 (2023).

31. Nakamura, T., Wang, X., Terashi, G. & Kihara, D. DAQ-Score Database: assessment of map–model compatibility for protein structure models from cryo-EM maps. Nat. Methods 20, 775–776 (2023).

32. Sun, K. et al. Predicting Ca2+ and Mg2+ ligand binding sites by deep neural network algorithm. BMC Bioinformatics 22, 324 (2022).

33. Metal3D: a general deep learning framework for accurate metal ion location prediction in proteins | Nature Communications. https://www.nature.com/articles/s41467-023-37870-6.

34. Mohamadi, A. et al. An ensemble 3D deep-learning model to predict protein metal-binding site. Cell Rep. Phys. Sci. 3, 101046 (2022).

35. Rogers, D. & Hahn, M. Extended-Connectivity Fingerprints. J. Chem. Inf. Model. 50, 742–754 (2010).

36. Capecchi, A., Probst, D. & Reymond, J.-L. One molecular fingerprint to rule them all: drugs, biomolecules, and the metabolome. J. Cheminformatics 12, 43 (2020).

37. Boldini, D., et al. Effectiveness of Molecular Fingerprints for Exploring the Chemical Space of Natural Products. https://chemrxiv.org/engage/chemrxiv/article-details/6582a1fd9138d23161181406 (2023) doi:10.26434/chemrxiv-2023-0m355-v2.

38. Willett, P. Similarity-based virtual screening using 2D fingerprints. Drug Discov. Today 11, 1046–1053 (2006).

39. Soares, T. A. et al. The (Re)-Evolution of Quantitative Structure–Activity Relationship (QSAR) Studies Propelled by the Surge of Machine Learning Methods. J. Chem. Inf. Model. 62, 5317–5320 (2022).

40. Wójcikowski, M., Kukiełka, M., Stepniewska-Dziubinska, M. M. & Siedlecki, P. Development of a protein-ligand extended connectivity (PLEC) fingerprint and its application for binding affinity predictions. Bioinforma. Oxf. Engl. 35, 1334–1341 (2019).

41. Da, C. & Kireev, D. Structural Protein–Ligand Interaction Fingerprints (SPLIF) for Structure-Based Virtual Screening: Method and Benchmark Study. J. Chem. Inf. Model. 54, 2555–2561 (2014).

42. Sánchez-Cruz, N., Medina-Franco, J. L., Mestres, J. & Barril, X. Extended connectivity interaction features: improving binding affinity prediction through chemical description. Bioinformatics 37, 1376–1382 (2021).

43. Gainza, P. et al. Deciphering interaction fingerprints from protein molecular surfaces using geometric deep learning. Nat. Methods 17, 184–192 (2020).

44. Lyu, J. et al. Ultra-large library docking for discovering new chemotypes. Nature 566, 224–229 (2019).

45. Brewerton, S. C. The use of protein-ligand interaction fingerprints in docking. Curr. Opin. Drug Discov. Devel. 11, 356–364 (2008).

46. Fassio, A. V. et al. Prioritizing Virtual Screening with Interpretable Interaction Fingerprints. J. Chem. Inf. Model. 62, 4300–4318 (2022).

47. Fassio, A. V. et al. Prioritizing Virtual Screening with Interpretable Interaction Fingerprints. J. Chem. Inf. Model. 62, 4300–4318 (2022).

48. Guillaumin, M., Verbeek, J. & Schmid, C. Is that you? Metric learning approaches for face identification. in 2009 IEEE 12th International Conference on Computer Vision 498–505 (IEEE, Kyoto, 2009). doi:10.1109/ICCV.2009.5459197.

49. Yilmaz, S. F. & Kozat, S. S. Unsupervised Anomaly Detection via Deep Metric Learning with End-to-End Optimization. Preprint at http://arxiv.org/abs/2005.05865 (2020).

50. Bromley, J. et al. Signature verification using a “siamese” time delay neural network. Int. J. Pattern Recognit. Artif. Intell. 07, 669–688 (1993).

51. Kaya, M. & Bilge, H. Deep Metric Learning: A Survey. Symmetry 11, 1066 (2019).

52. Coupry, D. E. & Pogány, P. Application of deep metric learning to molecular graph similarity. J. Cheminformatics 14, 11 (2022).

53. Wu, F., Courty, N., Jin, S. & Li, S. Z. Improving molecular representation learning with metric learning-enhanced optimal transport. Patterns 4, 100714 (2023).

54. Ge, W., Huang, W., Dong, D. & Scott, M. R. Deep Metric Learning with Hierarchical Triplet Loss. Preprint at 10.48550/arXiv.1810.06951 (2018).

55. Hoffer, E. & Ailon, N. Deep metric learning using Triplet network. Preprint at http://arxiv.org/abs/1412.6622 (2018).

56. Ding, S., Lin, L., Wang, G. & Chao, H. Deep Feature Learning with Relative Distance Comparison for Person Re-identification. Preprint at http://arxiv.org/abs/1512.03622 (2015).

57. Kokhlikyan, N. et al. Captum: A unified and generic model interpretability library for PyTorch. Preprint at http://arxiv.org/abs/2009.07896 (2020).

58. Sundararajan, M., Taly, A. & Yan, Q. Axiomatic Attribution for Deep Networks. Preprint at http://arxiv.org/abs/1703.01365 (2017).

59. Laitaoja, M., Valjakka, J. & Jänis, J. Zinc coordination spheres in protein structures. Inorg. Chem. 52, 10983–10991 (2013).

60. Piovesan, D., Profiti, G., Martelli, P. L. & Casadio, R. The human “magnesome”: detecting magnesium binding sites on human proteins. BMC Bioinformatics 13 Suppl 14, S10 (2012).

61. Kirberger, M. et al. Statistical analysis of structural characteristics of protein Ca2+-binding sites. J. Biol. Inorg. Chem. JBIC Publ. Soc. Biol. Inorg. Chem. 13, 1169–1181 (2008).

62. Israeli, H. et al. Structure reveals the activation mechanism of the MC4 receptor to initiate satiation signaling. Science 372, 808–814 (2021).

63. Heyder, N. A. et al. Structures of active melanocortin-4 receptor–Gs-protein complexes with NDP-α-MSH and setmelanotide. Cell Res. 31, 1176–1189 (2021).

64. Zhang, H. et al. Structural insights into ligand recognition and activation of the melanocortin-4 receptor. Cell Res. 31, 1163–1175 (2021).

65. Yu, J. et al. Determination of the melanocortin-4 receptor structure identifies Ca2+ as a cofactor for ligand binding. Science 368, 428–433 (2020).

66. Yip, K. M., Fischer, N., Paknia, E., Chari, A. & Stark, H. Atomic-resolution protein structure determination by cryo-EM. Nature 587, 157–161 (2020).

67. Maki-Yonekura, S., Kawakami, K., Takaba, K., Hamaguchi, T. & Yonekura, K. Measurement of charges and chemical bonding in a cryo-EM structure. Commun. Chem. 6, 1–8 (2023).

68. Nakane, T. et al. Single-particle cryo-EM at atomic resolution. Nature 587, 152–156 (2020).

69. Zhang, K., Pintilie, G. D., Li, S., Schmid, M. F. & Chiu, W. Resolving individual atoms of protein complex by cryo-electron microscopy. Cell Res. 30, 1136–1139 (2020).

70. Zheng, W. et al. The Crystal Structure of Human Isopentenyl Diphosphate Isomerase at 1.7 Å Resolution Reveals its Catalytic Mechanism in Isoprenoid Biosynthesis. J. Mol. Biol. 366, 1447–1458 (2007).

71. Duvenaud, D. et al. Convolutional Networks on Graphs for Learning Molecular Fingerprints. Preprint at http://arxiv.org/abs/1509.09292 (2015).

72. Gale-Day, Z. J., Shub, L., Chuang, K. V. & Keiser, M. J. Proximity Graph Networks: Predicting Ligand Affinity with Message Passing Neural Networks. Preprint at 10.26434/chemrxiv-2024-hznxh (2024).

73. Heid, E. et al. Chemprop: A Machine Learning Package for Chemical Property Prediction. J. Chem. Inf. Model. 64, 9–17 (2024).

74. Maggiora, G., Vogt, M., Stumpfe, D. & Bajorath, J. Molecular Similarity in Medicinal Chemistry. J. Med. Chem. 57, 3186–3204 (2014).

75. Weininger, D., Weininger, A. & Weininger, J. L. SMILES. 2. Algorithm for generation of unique SMILES notation. J. Chem. Inf. Comput. Sci. 29, 97–101 (1989).

76. Pedregosa, F. et al. Scikit-learn: Machine Learning in Python. J. Mach. Learn. Res. JMLR.

77. Musgrave, K., Belongie, S. & Lim, S.-N. PyTorch Metric Learning. Preprint at 10.48550/arXiv.2008.09164 (2020).

78. Paszke, A. et al. PyTorch: An Imperative Style, High-Performance Deep Learning Library. Preprint at http://arxiv.org/abs/1912.01703 (2019).

79. Akiba, T., Sano, S., Yanase, T., Ohta, T. & Koyama, M. Optuna: A Next-generation Hyperparameter Optimization Framework. Preprint at http://arxiv.org/abs/1907.10902 (2019).

80. Virtanen, P. et al. SciPy 1.0: fundamental algorithms for scientific computing in Python. Nat. Methods 17, 261–272 (2020).

