## SupplementalTables1-7 for "Metric Ion Classification (MIC): A deep learning tool for assigning ions and waters in cryo-EM and x-ray crystallography structures": SIGuide.doc.docx

SI Guide

Supplementary Table 1. (external Excel file)

Selected PDB sites used in training and testing the MIC metric embedding models. Third column contains the deposited ion identity, used as the ground-truth label during training. Fifth column indicates if the example was used for training or held-out for testing. Sixth column specifies if a given example was in the prevalent or extended set.

Supplementary Table 2. (external Excel file)

MIC predictions and manual annotations on the test set, generated using the limited-set model. The fifth column shows the MIC-predicted class. Columns seven through twelve show the prediction of each output class for a given example, in order: water, magnesium, sodium, zinc, calcium, chlorine.

Supplementary Table 3. (external Excel file)

Features corresponding to the top ten bit features by attribution with integrated gradients for the representative calcium-zinc and magnesium-zinc pairs. Column one is the index in the fingerprint assigned the attribution, column two is the calculated attribution value, column 3 is the bit value that is folded to the corresponding fingerprint index; note that due to bit collisions, multiple bits can hash to the same index. Columns four and five contain the shell level and radius, respectively. Column six is the central atom group of the shell. Column seven is the number of interactions in a shell, and column eight is whether the attribution was calculated against magnesium or calcium.

Supplementary Table 4. (external Excel file)

MIC outputs for the Cryo-EM MC4R and apoferritin ions. The first four columns are the PDB ID, chain, residue index, and deposited label. Columns five through ten show the prediction of each output class for a given example, in order: water, magnesium, sodium, zinc, calcium, chlorine. Column eleven is the predicted class, and column 12 is the model confidence. Column 12 specifies if the structure is MC4R or apoferritin.

Supplementary Table 5. (external Excel file)

RNA/Ribosome full sites and predictions, prevalent-only MIC model. Columns one through four are the PDB ID, chain, ion index, and residue name in the file deposited in the PDB. Columns five through ten are the individual MIC-predicted probabilities for each class in the prevalent set: water, magnesium, sodium, zinc, calcium, chloride. Columns eleven and twelve are the MIC prediction and confidence.

Supplementary Table 6. (external Excel file)

Test set predictions and annotations of the extended-set MIC model. Columns one through four are the PDB ID, chain, deposited residue name, and residue index in the PDB of each test example. Columns five and six are the MIC prediction and confidence. Columns seven through seventeen are the MIC output probabilities for each class in the extended set: water, magnesium, sodium, zinc, calcium, chloride, potassium, manganese, iodide, iron, and bromide.

Supplementary Table 7. (external Excel file)

MIC prevalent-model and Undowser cumulative clash predictions on a selection of PDB structures with all ions renamed to waters. Columns one through four are the PDB id, chain, residue name, and residue index of each ion and water in the evaluated structures. Columns five and six are the MIC predictions and confidence, and columns seven through twelve are the individual predictions for the six classes in the prevalent-set model: water, magnesium, sodium, zinc, calcium, and chlorine. Column twelve is whether a given ion or water was flagged by Undowser as likely not being a water. Column thirteen is the cumulative severity of the clashes as output by Undowser.
